# Functional remodeling of the parasubthalamic nucleus drives alcohol drinking escalation in dependence

**DOI:** 10.64898/2026.04.30.722112

**Authors:** Jeffery L Dunning, Max Kreifeldt, Agbonlahor Okhuarobo, Catherine Lopez, Florence P Varodayan, Michal Bajo, Rachel J Smith, Celeste Moreau, Colton Krull, Leeann Shu, Allison White, Giovana Macedo, Harpreet Sidhu, Howard C Becker, Marisa Roberto, Karl Deisseroth, Candice Contet

## Abstract

The neurocircuitry changes mediating the development and maintenance of an alcohol use disorder are complex and dynamic. The parasubthalamic nucleus (PSTN), a small nucleus of the posterior lateral hypothalamus best known for suppressing appetite, is interconnected with brain regions disrupted in addiction; yet its potential role in the regulation of alcohol consumption had never been examined. Here we show that the PSTN exerts potent control over alcohol drinking in mice. Remarkably, the influence of endogenous PSTN activity on voluntary alcohol consumption switches from inhibitory to stimulatory upon induction of alcohol dependence. Among PSTN cells, *Crh* neurons represent a unique subpopulation that promotes alcohol drinking and fires more in dependent mice. Alcohol intake escalation driven by PSTN *Crh* neurons involves thalamic output and behavioral disinhibition. Based on our results, PSTN *Crh* neurons could represent a critical node in the brain circuitry overactive in alcohol addiction driven by reward seeking in humans.

## Introduction

The development and maintenance of an alcohol use disorder (AUD) result from neural dysfunction in three domains: incentive salience, negative emotionality, and executive control ^1,2^. Whether these three factors are intercorrelated (i.e., dynamically at play in the same individual) or independent from each other (i.e., defining mechanistic subtypes of alcohol misuse, each predominantly driven by one factor) is a matter of debate ^3,4^. The multidimensional and heterogeneous etiology of alcohol addiction has critical implications for treatment - a medication targeting dysfunction in one domain will have differential efficacy depending on the relative contribution of that domain to alcohol misuse in each patient at a given time. Functional neuroimaging in humans and mechanistic studies in rodents have identified brain regions associated with each domain whose activity confers vulnerability to AUD and/or is altered by chronic alcohol exposure ^1,4–6^.

The parasubthalamic nucleus (PSTN) is a small nucleus of the posterior lateral hypothalamus that is highly interconnected with some of these brain regions ^7^. Notably, the PSTN sends strong projections to elements of the “extended amygdala”, a macrostructure of the basal forebrain whose recruitment during the withdrawal phase of the addiction cycle is thought to produce negative emotionality and drive compulsive alcohol drinking through negative reinforcement ^1,8–10^. Moreover, the PSTN contains a cluster of cells expressing the neuropeptide corticotropin-releasing factor (CRF, encoded by the *Crh* gene) and CRF signaling in extended amygdala subregions is known to promote alcohol consumption and hyperkatifeia (e.g., negative affect, increased pain sensitivity) in animal models of alcohol dependence ^10–14^. Therefore, we hypothesized that the PSTN might serve as a critical source of CRF recruited in alcohol dependence.

The PSTN is receiving increasing attention for its role in appetite suppression and stress responses ^15–27^, but it has never been examined in the context of addiction. We therefore sought to establish the role of PSTN activity in the control of alcohol drinking and to elucidate the underlying signaling, circuit, and motivational mechanisms, with a particular interest in the *Crh* subpopulation of PSTN neurons.

## Results

### The influence of PSTN activity on alcohol drinking switches from inhibitory to stimulatory upon induction of alcohol dependence

We first determined whether PSTN activity is sensitive to chronic intermittent ethanol (CIE) vapor inhalation, a modality of alcohol exposure that mimics the repeated cycles of prolonged intoxication and withdrawal experienced by individuals with an AUD. Exposing mice to CIE has translational relevance for AUD as it gradually escalates their voluntary alcohol consumption to intoxicating levels when combined with limited-access, free-choice drinking sessions (2-h alcohol/water two-bottle choice [2BC]) on alternating weeks ^28,29^ (Fig. 1A). After five rounds of CIE/2BC alternation, we found that PSTN activity (indexed by expression of the immediate early gene product c-Fos) was blunted during vapor inhalation (0 h withdrawal timepoint), rebounded in early withdrawal (8 h and 24 h), and returned to control levels after 72 h when comparing CIE-exposed male mice to counterparts breathing air only (Fig. 1B-C). Normalization of PSTN cFos+ cell counts at 7 days post-vapor was independent of resuming alcohol drinking sessions in the last week vs. keeping the mice abstinent (Fig. 1C). The same pattern of PSTN silencing during intoxication followed by transient activation during early withdrawal was observed in mice exposed to CIE without 2BC experience (Fig. S1A).

**Figure 1.**
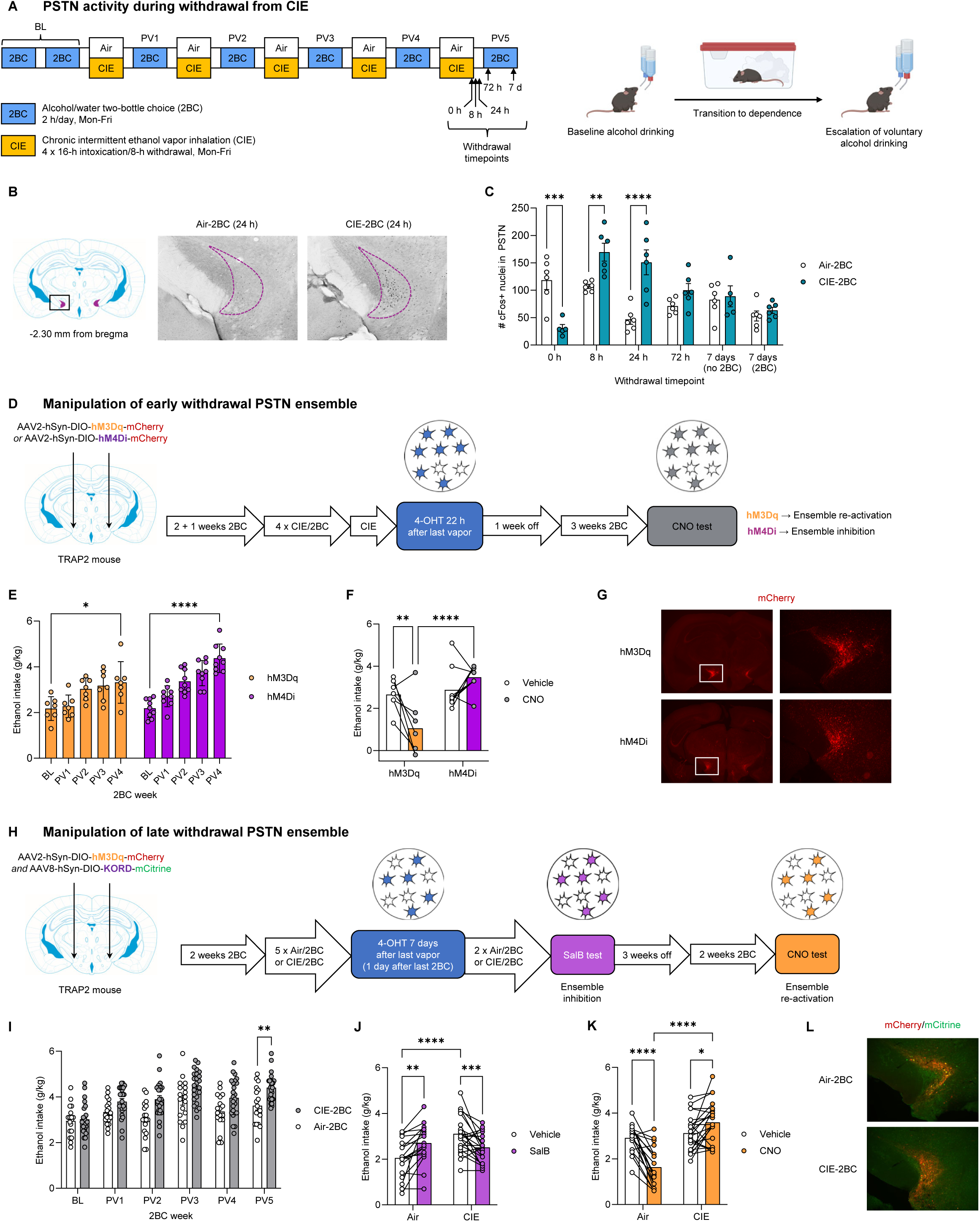
The influence of PSTN activity on alcohol drinking switches from inhibitory to stimulatory upon induction of alcohol dependence. A. Experimental strategy for the measure of neuronal activation in the PSTN during withdrawal following a history of voluntary alcohol drinking combined, or not, with chronic intermittent alcohol vapor inhalation. BL, baseline; PV, post-vapor. B. Representative images of c-Fos immunoreactivity in the PSTN during early withdrawal. C. Alcohol intoxication reduced, while early withdrawal increased the number of c-Fos+ cells in the PSTN. Air-2BC vs. CIE-2BC: **, p<0.01; ***, p<0.001; ****, p<0.0001. D. Experimental strategy for the expression of designer receptors in PSTN neurons active in early withdrawal from CIE and their subsequent chemogenetic re-activation or inhibition. E. Alcohol intake escalation prior to 4-OHT administration. BL vs. PV4: *, p<0.05; ****, p<0.0001. F. Re-activating the early withdrawal PSTN ensemble reduced alcohol intake. Pairwise comparisons: **, p<0.01; ****, p<0.0001. G. Labeling of the early withdrawal PSTN ensemble by mCherry. H. Experimental strategy for the expression of designer receptors in PSTN neurons active in late withdrawal from CIE and their subsequent chemogenetic re-activation or inhibition. I. Alcohol intake escalation prior to 4-OHT administration. Air-2BC vs. CIE-2BC: **, p<0.01. J. Inhibiting the late withdrawal PSTN ensemble increased alcohol intake in Air-2BC but reduced it in CIE-2BC mice. Pairwise comparisons: **, p<0.01; ***, p<0.001; ****, p<0.0001. K. Re-activating the late withdrawal PSTN ensemble reduced alcohol intake in Air-2BC but increased it in CIE-2BC mice. Pairwise comparisons: *, p<0.05; ****, p<0.0001. L. Labeling of the late withdrawal PSTN ensemble by mCherry and mCitrine. See also Figure S1 and Table S2.

To determine the functional significance of PSTN activity for alcohol drinking, we prepared TRAP2 mice, which express an inducible Cre recombinase in the *Fos* locus, to trigger the expression of designer receptors (hM3Dq, hM4Di, or KORD) in active PSTN cells upon administration of 4-hydroxytamoxifen (4-OHT). This strategy enables subsequent chemogenetic re-activation (upon activation of hM3Dq by clozapine-N-oxide [CNO]) or inhibition (upon activation of hM4Di by CNO or KORD by salvinorin B [SalB]) of the targeted ensemble ^30–33^. We verified that no recombination occurs in the PSTN of TRAP2 mice unless they receive 4-OHT (Fig. S1B, see also ^21^).

A cohort of TRAP2 male and female mice prepared for PSTN expression of hM3Dq or hM4Di was exposed to five rounds of CIE/2BC alternation (Fig. 1D), which significantly increased their alcohol drinking compared to pre-CIE baseline levels (Fig. 1E, S1C). They were injected with 4-OHT 22 h after vapor inhalation ended to target the early withdrawal ensemble. Re-activating the early withdrawal PSTN ensemble after four weeks of vapor withdrawal reduced alcohol intake while inhibiting it had no significant effect (Fig. 1F-G, S1D). This result indicates that the PSTN cells active during early withdrawal from severe intoxication suppress alcohol intake but do not remain active during protracted withdrawal.

Importantly, in the CIE-2BC model, alcohol drinking ramps up during late (3-7 days) rather than early withdrawal from vapor ^28,34^. Although the number of cFos+ PSTN cells were similar between Air and CIE mice at the 72-h and 7-day timepoints (Fig. 1C), we reasoned that these two ensembles may be made up of different PSTN cells, which may not exert the same influence on alcohol drinking. To test this hypothesis, another mixed-sex cohort of TRAP2 mice was prepared for within-subject PSTN expression of hM3Dq and KORD, exposed to five rounds of CIE (or air)/2BC alternation, and injected with 4-OHT 7 days after vapor, 22 h after 2BC (Fig. 1H). As expected, CIE mice consumed higher levels of alcohol than Air mice prior to 4-OHT administration (Fig. 1I), albeit this difference reached significance only in males (Fig. S1E-F,I-J). 4-OHT was administered at the usual time of 2BC start but the mice were not given access to alcohol that day, thus capturing PSTN activity associated with the motivation to drink alcohol rather than the effect of alcohol itself. The mice were exposed to two more rounds of CIE/2BC alternation to maintain alcohol intake escalation while allowing full expression of the viral constructs. Inhibiting the PSTN ensembles had opposite effects in the two groups: it increased voluntary alcohol drinking in Air mice but reduced it in CIE mice, thus ablating the intake difference between the two groups (Fig. 1J). Five weeks later, without further exposure to CIE,

Air and CIE mice consumed similar levels of alcohol and re-activating their respective PSTN ensembles again produced opposite effects: it reduced alcohol intake in Air mice but increased it in CIE mice, thus reinstating the group difference that was present at the time of Cre induction (Fig. 1K-L). These effects were seen in both males and females (Fig. S1G-H,K-L). These data demonstrates that the PSTN undergoes drastic functional remodeling upon exposure to CIE, such that the influence of endogenous PSTN activity on alcohol drinking switches from inhibitory in non-dependent drinkers to stimulatory in dependent drinkers. We reasoned that this outcome reflects the recruitment of a PSTN subpopulation promoting alcohol intake, which neutralizes the suppression exerted by other PSTN neurons.

### PSTN *Crh* neurons promote alcohol drinking and are more excitable in alcohol dependence

To investigate the contribution of *Crh* neurons to the influence of PSTN ensembles on alcohol drinking, *Crh*-Cre mice were prepared to express hM4Di in PSTN *Crh* cells and exposed to five rounds of CIE (or air)/2BC alternation. Male mice were used for this experiment as CIE-induced alcohol intake escalation is typically more robust in this sex (^35–39^ and Fig. S1E,I). As expected, CIE mice consumed higher levels of alcohol than Air counterparts (Fig. 2B, S2A). Inhibiting PSTN *Crh* cells reduced alcohol drinking in both groups and CNO-treated CIE mice reduced their intake to the level of vehicle-treated Air mice (Fig. 2C). This result indicates that the endogenous activity of PSTN *Crh* neurons promotes alcohol consumption in both non-dependent and dependent drinkers.

**Figure 2.**
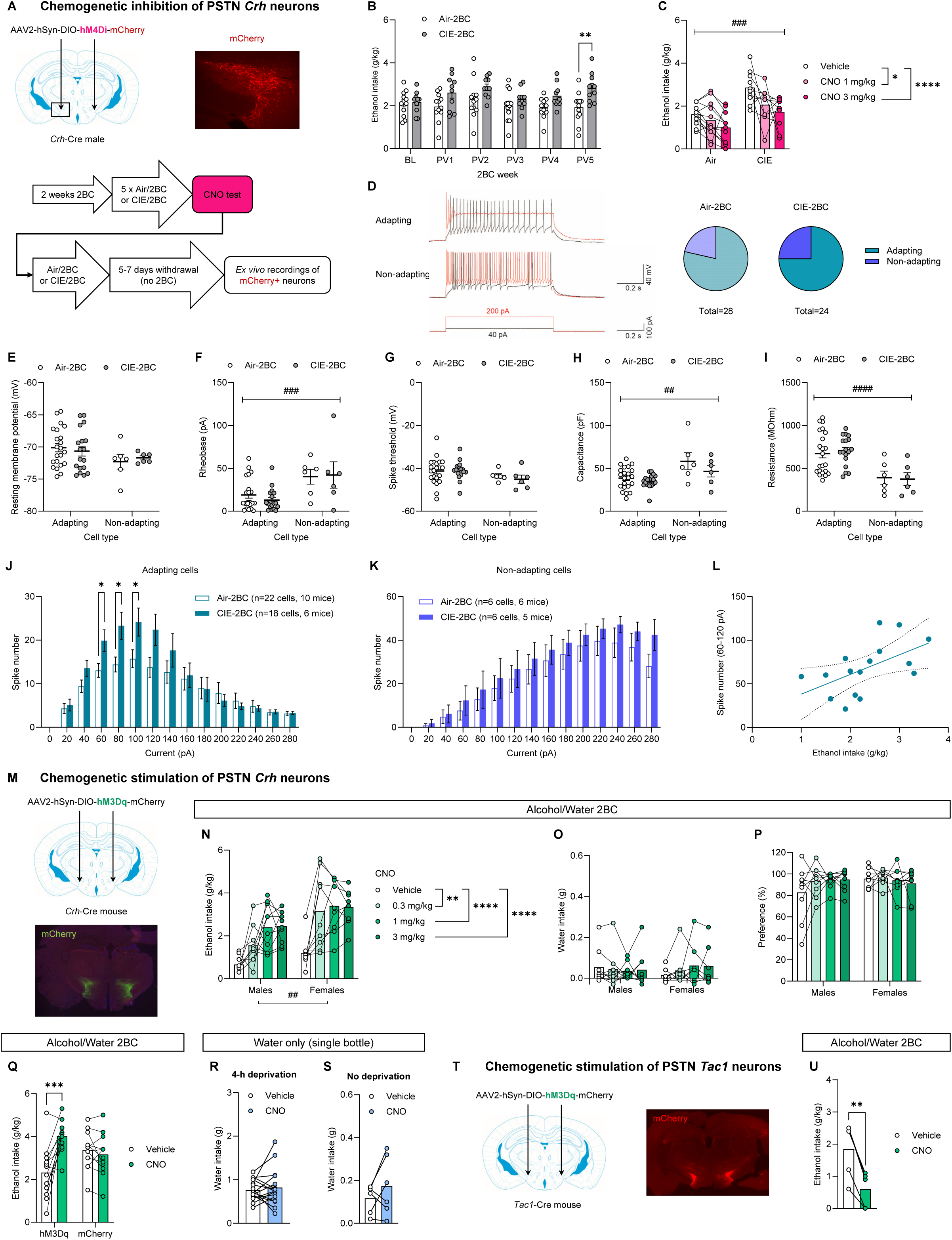
PSTN *Crh* neurons promote alcohol drinking and are more excitable in alcohol dependence. A. Experimental strategy for chemogenetic inhibition and electrophysiological recordings of PSTN *Crh* neurons during late withdrawal from CIE. B. Alcohol intake escalation prior to CNO testing. Air-2BC vs. CIE-2BC: **, p<0.01. C. Inhibiting PSTN *Crh* neurons reduced alcohol drinking. Main effect of vapor: ###, p<0.001. Comparisons to Vehicle: *, p<0.05; ****, p<0.0001. D. Representative traces of adapting (left) and non-adapting (right) PSTN *Crh* cells during 40 pA (black) and 200 pA (red) sweeps. The proportion of each cell type was equivalent in Air-2BC and CIE-2BC mice. E-I. Membrane properties of adapting and non-adapting PSTN *Crh* cells: resting membrane potential (E), rheobase (F), spike threshold (G), capacitance (H), and resistance (I). Main effect of cell type: ##, p<0.01; ###, p<0.001; ####, p<0.001. J-K. CIE increased the number of spikes evoked by 60-100 pA currents in mCherry-labeled PSTN *Crh* adapting cells (J) but had no effect in non-adapting cells (K). Air-2BC vs. CIE-2BC: *, p<0.05. L. Positive correlation between spiking in adapting cells and recent alcohol intake, p<0.05. M. Experimental strategy for chemogenetic activation of PSTN *Crh* neurons. N-P. Activating PSTN *Crh* neurons increased alcohol intake (N) without affecting water intake (O) or alcohol preference (P). Comparisons to Vehicle: **, p<0.01; ****, p<0.0001. Main effect of sex: ##, p<0.01. Q. The effect of CNO on alcohol intake requires hM3Dq expression. Vehicle vs. CNO: ***, p<0.001. R-S. CNO had no effect on water intake in deprived (R) or quenched (S) states. T. Experimental strategy for chemogenetic activation of PSTN *Tac1* neurons. U. Activating PSTN *Tac1* neurons reduced alcohol intake. Vehicle vs. CNO: **, p<0.01. See also Figure S2 and Table S2.

To evaluate whether PSTN *Crh* neurons may be recruited by alcohol dependence, the same mice were exposed to additional rounds of CIE/2BC and the excitability of PSTN *Crh* cells was measured via slice electrophysiology 5-7 days after last vapor, using the mCherry reporter fused to hM4Di to identify them (Fig. 2A). We verified that CNO inhibited firing in mCherry+ cells (Fig. S2B) while firing rates remained stable in the absence of CNO (rundown control, Fig. S2C). The input-output curves identified two populations of PSTN *Crh* neurons: most cells (henceforth designated as “adapting”) displayed an inverted U-curve upon injection of increasing currents with a peak ∼100 pA, while a minority of cells (henceforth designated as “non-adapting”) increased their firing rates up to current steps of 200 pA and higher (Fig. 2D,J-K). Non-adapting cells had significantly higher rheobase, higher capacitance, and lower resistance than adapting cells, but CIE exposure did not affect any of these parameters in either population (Fig. 2E-I). In contrast, the firing rate of adapting cells was significantly higher in CIE mice than in Air counterparts at current steps nearing the peak (Fig. 2J), while no group difference was detected for non-adapting cells (Fig. 2K). Further supporting a functional link between the excitability of PSTN *Crh* adapting cells and alcohol dependence, the cumulative number of spikes fired by adapting cells in the 60-120 pA range was positively correlated with ethanol intake during the last 2BC week (Fig. 2L). These results show that a subset of PSTN *Crh* cells becomes more excitable upon CIE exposure and that their increased activity contributes to alcohol drinking escalation in dependent mice.

Next, we leveraged chemogenetic stimulation to mimic the effect of CIE. From that point onwards, all experiments used mixed-sex cohorts. *Crh*-Cre mice were prepared for hM3Dq expression in the PSTN (Fig. 2M). We used cFos immunohistochemistry to verify that CNO activated PSTN cells labeled with the mCherry reporter fused to hM3Dq (Fig. S2D-F).

Stimulating PSTN *Crh* neurons produced a robust increase in alcohol drinking across sexes (Fig. 2N) without altering water intake (negligible in this limited-access model) or alcohol preference (Fig. 2O-P). As expected, CNO did not affect alcohol intake in mice expressing the mCherry reporter alone, confirming that the effects of CNO are mediated by hM3Dq (Fig. 2Q). Furthermore, stimulating PSTN *Crh* neurons did not affect water intake in quenched or deprived states, thus indicating that the increased consumption of alcohol does not result from thirst (Fig. 2R-S). Stimulating PSTN *Crh* neurons via optogenetics also resulted in increased alcohol drinking (Fig. S2G-I).

The influence of PSTN *Crh* neurons on alcohol intake contrasts with the well-established role of PSTN activity in feeding and drinking suppression ^7,16,17,21,24^. In particular, the PSTN contains a large population of neurons expressing substance P (encoded by *Tac1*) that shows little overlap with the *Crh* population and whose chemogenetic or optogenetic stimulation reduces food and sucrose consumption ^17^. To verify that the phenotypes we observed in *Crh*-Cre mice did not result from technical idiosyncrasies, we tested *Tac1*-Cre mice under the same experimental conditions (Fig. 2T). As expected, stimulating PSTN *Tac1* neurons via hM3Dq strongly reduced alcohol intake (Fig. 2U). Altogether, these results highlight that *Crh* neurons represent a unique subpopulation within the PSTN that promotes alcohol consumption, thus opposing the influence of *Tac1* neurons.

### Alcohol consumption driven by PSTN *Crh* neurons involves glutamate and neuropeptide signaling

We next sought to address the signaling mechanism that mediates the increase in alcohol drinking driven by PSTN *Crh* neurons. Given the literature implicating CRF_1_ signaling in excessive alcohol drinking ^10,11,13,40^, we reasoned that blocking CRF_1_ receptors would prevent this effect. *Crh*-Cre mice expressing hM3Dq in the PSTN and trained to drink alcohol in 2-h free choice sessions were co-injected intraperitoneally (i.p.) with CNO (or vehicle) and different doses of the CRF_1_ receptor antagonist CP376395 before 2BC. CP376395 tended to reduce alcohol intake in vehicle-injected mice but alcohol consumption following CNO administration was insensitive to CRF_1_ blockade (Fig. 3A, S3A). This result was replicated in a separate cohort of mice, in which intake was also measured after a shorter period (1 h) of alcohol access (Fig. S3B-D). To test whether CRF signaling at other targets (e.g., CRF_2_ receptors) might be implicated, we blocked CRF synthesis in the PSTN using a short hairpin RNA (shRNA) targeted against *Crh* (shCrh). We previously determined that the shCrh vector reduces PSTN *Crh* expression by >90% compared to a vector encoding a control shRNA sequence (shControl) ^41^.

**Figure 3.**
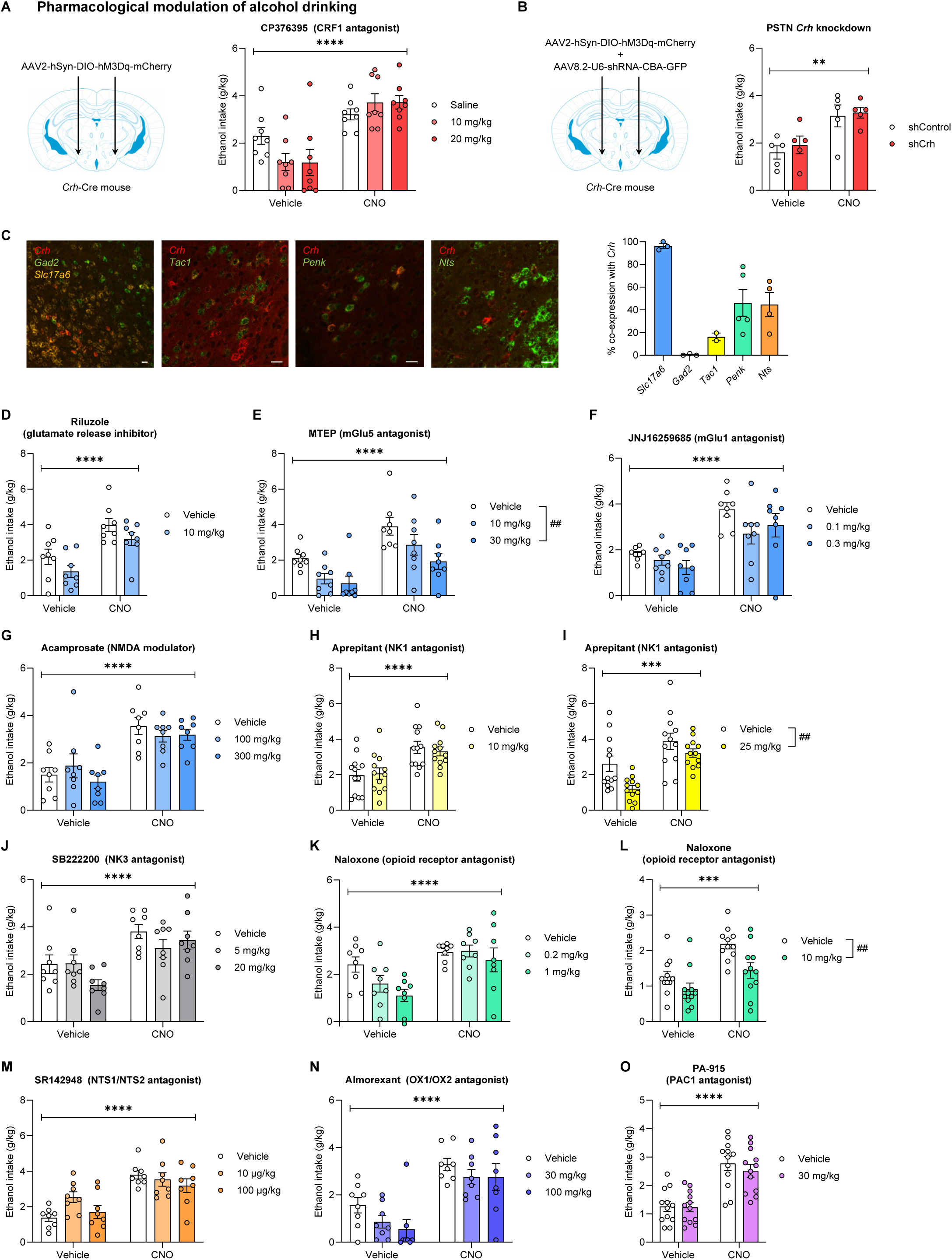
Alcohol consumption driven by PSTN *Crh* neurons involves glutamate and neuropeptide signaling. A. *Crh*-Cre mice were injected in the PSTN with a Cre-dependent hM3Dq-encoding vector. A CRF_1_ antagonist (CP376395) was administered at the same time as CNO 30 min prior to alcohol 2BC. Ethanol intake was measured following combined administration of CP376395 (between-subjects) and CNO (within-subjects). Main effect of CNO: ****, p<0.0001. B. *Crh*-Cre mice were injected in the PSTN with a Cre-dependent hM3Dq-encoding vector, along with a vector encoding an shRNA targeting *Crh* (shCrh) or a control shRNA sequence (shControl). CNO was administered 30 min prior to alcohol 2BC. Main effect of CNO: **, p<0.01. C. The cellular colocalization of *Crh* mRNA with markers of GABAergic (*Gad2*) or glutamatergic (*Slc17a6*) neurons, or with neuropeptide-encoding mRNAs known to be expressed in the PSTN (*Tac1*, *Penk*, *Nts*), was visualized by fluorescent *in situ* hybridization. D-O. Mice were prepared as in A. A ligand (D-K, M-N: between-subjects, **L,O**: within-subjects) was administered at the same time as CNO (within-subjects) 30 min prior to alcohol 2BC. Main effect of CNO: ***, p<0.001; ****, p<0.0001. Dunnett’s *posthoc* test vs. Vehicle (E) or main effect of ligand (I,L): ^##^, p<0.01. See also Figure S3 and Table S2.

Stimulating PSTN *Crh* neurons increased alcohol intake to the same extent in shControl and shCrh mice (Fig. 3B). Altogether, these findings indicate that the increased alcohol drinking produced by chemogenetic stimulation of PSTN *Crh* neurons does not require CRF signaling. We reasoned that other neurotransmitters or neuromodulators released by PSTN *Crh* neurons might contribute to the effect of their chemogenetic stimulation on alcohol drinking. We characterized the co-localization of *Crh* with markers of glutamatergic (*Scl17a6*) and GABAergic (*Gad2*) neurons, as well as neuropeptide-encoding transcripts expressed at high levels in the PSTN (*Tac1*, *Penk*, *Nts*) (Fig. 3C). Consistent with the neurochemical makeup of the PSTN ^7^, virtually all PSTN *Crh* neurons express *Slc17a6* and not *Gad2*. Consistent with a previous report ^17^, we observed a limited overlap (∼15%) between *Crh* and *Tac1*. A larger fraction of PSTN *Crh* cells (∼40%) co-express *Penk* or *Nts*, the transcripts encoding enkephalins and neurotensin, respectively.

Following the same approach as with CP376395, we then tested whether antagonizing glutamate (glutamate release inhibitor riluzole, Fig. 3D; metabotropic glutamate receptor 5 (mGlu5) antagonist MTEP, Fig. 3E; mGlu1 antagonist JNJ16259685, Fig. 3F; NMDA receptor modulator acamprosate, Fig. 3G), substance P (neurokinin receptor 1 (NK1) antagonist aprepitant, Fig. 3H-I; NK3 antagonist SB222200, Fig. 3J), enkephalin (opioid receptor antagonist naloxone, Fig. 3K-L), or neurotensin (NTS1/NTS2 receptor antagonist SR142948, Fig. 3M) signaling would compromise the ability of CNO to increase voluntary alcohol drinking in *Crh*-Cre mice expressing hM3Dq in the PSTN. We also tested the potential involvement of orexin signaling (OX1/OX2 receptor antagonist almorexant, Fig. 3N), as this neuropeptide is produced in the lateral hypothalamic area, adjacent to the PSTN, where *Crh* neurons also reside. Partial co-localization (∼20%) of CRF and pituitary adenylate cyclase-activating polypeptide (PACAP) in the PSTN was recently reported ^19^, so we also tested the role of PACAP signaling (PAC_1_ receptor antagonist PA-915, Fig. 3O). For each target, mice were split into subgroups of equivalent baseline ethanol intake, each assigned to a given antagonist dose (Fig. S3E-M) for within-subject testing of CNO vs. vehicle (co-injected with the antagonist prior to alcohol 2BC). None of the ligands tested selectively blocked the effect of CNO, consistent with our observation that the endogenous activity of PSTN *Crh* neurons contributes to alcohol consumption even in moderate drinkers (Fig. 2C). As with CP376395, we found that SB222200, naloxone (0.2-1 mg/kg), and almorexant tended to reduce alcohol consumption in the vehicle condition but failed to exert the same effect in the CNO condition (Fig. 3J,K,N). On the other hand, MTEP, aprepitant (25 mg/kg), and naloxone (10 mg/kg) reduced alcohol drinking regardless of CNO vs. vehicle pretreatment (Fig. 3E,I,L). Alcohol intake following CNO injection remained higher than following vehicle injection, indicating that increasing the activity of PSTN *Crh* neurons can overcome the suppressive effect of mGlu5, opioid receptor, and NK1 blockade on alcohol drinking. In conclusion, alcohol drinking driven by PSTN *Crh* neurons involves a combination of glutamate and neuropeptide signaling and elevating their activity confers resistance to pharmacological manipulations that reduce alcohol drinking under control conditions.

### The influence of PSTN *Crh* neurons on alcohol consumption is pathway-dependent and PVT projectors drive escalation in dependent drinkers

To identify a neural pathway that might mediate the increase in alcohol drinking driven by PSTN *Crh* neurons, we first mapped out their projections using a viral tracer (Fig. 4A). Prominent tdTomato (axonal) and EGFP (presynaptic) labeling was observed across a continuum of basal forebrain structures (Fig. 4B-F), including elements of the extended amygdala, such as the interstitial nucleus of the posterior limb of the anterior commissure (IPAC, Fig. 4C-E, Fig. S4G) and sublenticular substantia innominata (labeled EA in the Paxinos atlas, Fig. 4D-E, Fig. S4E,G), and to a lesser extent, the ventral part of the BNST (Fig. 4C, Fig. S4E) and medial division of the CeA (Fig. 4E-F, Fig. S4G). The ventral pallidum (VP) was also densely labeled with both markers, especially at posterior levels, where the VP adjoins the IPAC (Fig. 4B-D, Fig. S4E). Several thalamic nuclei showed moderate innervation, including the PVT and its ventral neighbor, the intermediodorsal thalamic nucleus (IMD) (Fig. 4F, Fig. S4F,H). Prominent labeling was also observed in midbrain and brainstem areas, most notably throughout the reticular formation, as well as in the zona incerta, ventral tegmental area, periaqueductal gray matter, dorsal raphe, and parabrachial nucleus (Fig. S4).

**Figure 4.**
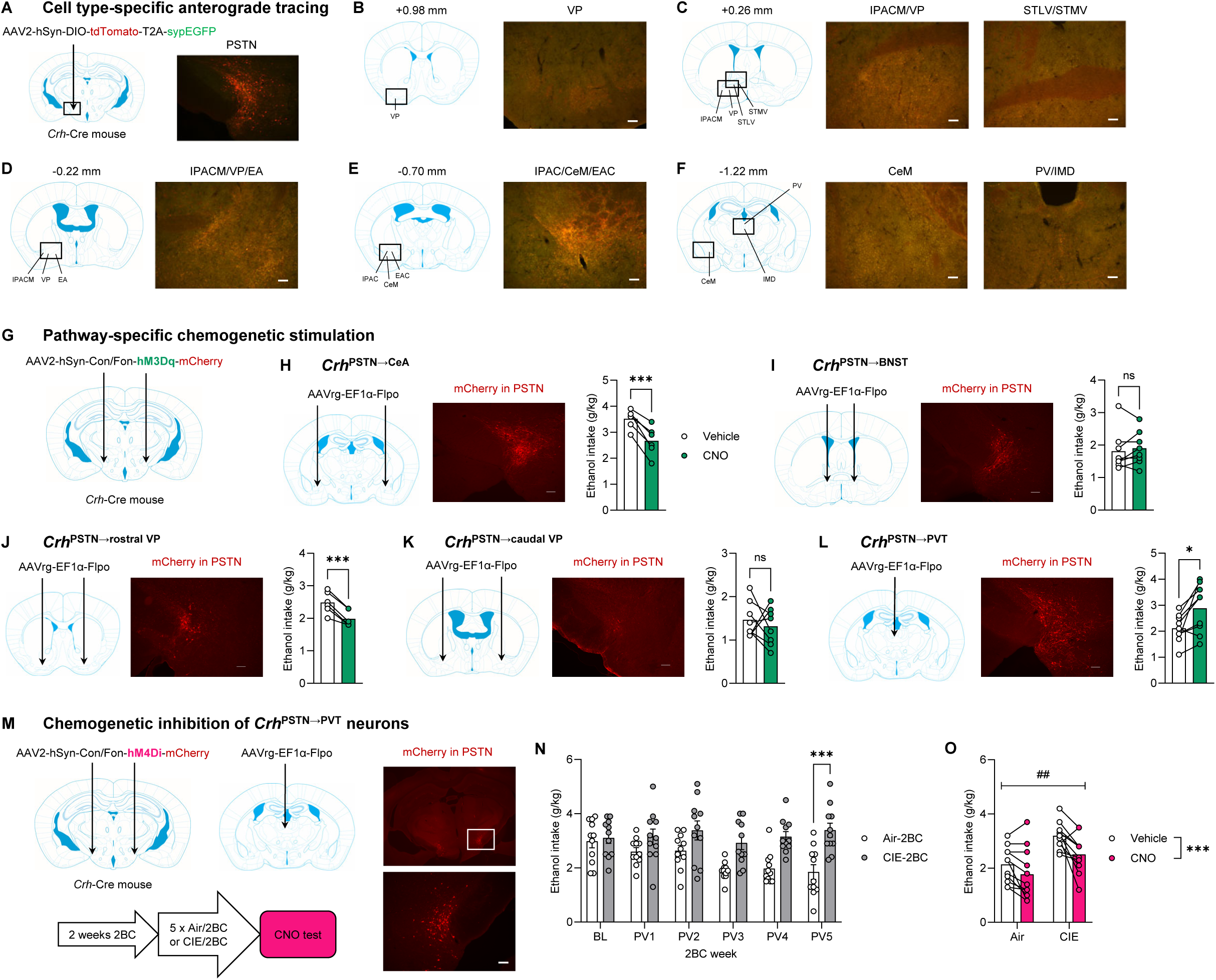
The influence of PSTN *Crh* neurons on alcohol consumption is pathway-dependent and PVT projectors drive escalation in dependent drinkers. A. *Crh*-Cre mice were injected unilaterally in the PSTN with a Cre-dependent vector encoding the green fluorescent protein EGFP fused to synaptophysin, a protein selectively targeted to presynaptic terminals, and the cytoplasmic red fluorescent protein tdTomato. B-F. EGFP and tdTomato immunolabeling (green and red respectively, overlaid) shows transduction in the PSTN (A) and innervation in basal forebrain (B-F) and thalamic (F) areas (scale bars, 100 µm). The field of view is outlined in the brain atlas diagram on the left; distance is anteroposterior from bregma; abbreviations from the Franklin and Paxinos atlas ^75^ are used (CeM, central amygdaloid nucleus, medial division; EA, extended amygdala; EAC, extended amygdala, central part; IMD, intermediodorsal thalamic nucleus; IPAC, interstitial nucleus of the posterior limb of the anterior commissure; IPACM, IPAC, medial part; PV, paraventricular thalamic nucleus; STMV, BNST, medial division, ventral part; STLV, BNST, lateral division, ventral part; VP, ventral pallidum). G. An intersectional strategy was used to express hM3Dq in PSTN *Crh* neurons projecting to specific forebrain targets. *Crh*-Cre mice were injected in a target region of interest with a retrograde vector encoding Flpo and in the PSTN with a Cre- and Flp-dependent (Con/Fon) vector encoding hM3Dq. H-L. The brain atlas diagram shows the site of injection of the retrograde vector (H, CeA; I, BNST; J, rostral VP; K, caudal VP; L, PVT) and the image shows mCherry immunolabeling in the PSTN (scale bars, 100 µm). Alcohol intake was measured following administration of CNO (within-subjects) 30 min prior to 2BC. CNO vs. Vehicle: *, p<0.05; ***, p<0.001, paired t-tests. See also Figure S4 and Table S2. M. The same intersectional strategy was used to express hM4Di in *Crh*^PSTN→PVT^ neurons, which are labeled with mCherry. CNO was tested during late withdrawal from CIE. N. Alcohol intake escalation prior to CNO testing. Air-2BC vs. CIE-2BC: ***, p<0.001. O. Inhibiting *Crh*^PSTN→PVT^ neurons reduced alcohol drinking. Main effect of vapor: ##, p<0.01. Main effect of CNO: ***, p<0.001.

We probed the functional role of individual forebrain projections using an intersectional viral strategy combining retrograde delivery of a Flpo-encoding vector in the target region with infusion of a Cre- and Flp-dependent (Con/Fon) hM3Dq vector in the PSTN of *Crh*-Cre mice ^21,42^ (Fig. 4G). Based on previous findings implicating CeA and BNST adaptations in AUD ^14,43^, we hypothesized that projections to one of these regions might mediate the increase in alcohol drinking driven by PSTN *Crh* neurons. In contrast to our predictions, activating CeA projectors reduced ethanol intake (Fig. 4H) while stimulating BNST (Fig. 4I) projectors had no significant effect. Turning to pallidal targets, we found that rostral VP projectors suppress ethanol intake (Fig. 4J) while caudal VP projectors exert no influence (Fig. 4K). In contrast, stimulating PVT projectors increased ethanol intake (Fig. 4L).

To determine whether PVT projectors are recruited in dependence, we used the same combinatorial strategy to express hM4Di in a pathway- and cell type-specific manner and exposed the mice to five rounds of CIE (or air)/2BC alternation (Fig. 4M). As expected, CIE mice consumed higher levels of alcohol than Air counterparts, an effect driven by males (Fig. 4N, S4S-T,V-W). Inhibiting *Crh*^PSTN→PVT^ neurons reduced alcohol drinking across groups, with CIE males being the most sensitive (Fig. 4O, S4U,X).

Altogether, these results reveal unexpected functional heterogeneity among PSTN *Crh* neurons, with PVT projectors replicating the influence of the general PSTN *Crh* population on alcohol drinking, an effect that is opposed by CeA and rostral VP projectors in non-dependent drinkers but dominates to drive intake escalation in dependent drinkers.

### Activating PSTN *Crh* neurons disinhibits behavior in response to novelty and stress

Increases in alcohol consumption can result from various sources of positive and negative reinforcement ^44,45^. We thus sought to determine the affective and nociceptive state of mice upon stimulation of PSTN *Crh* neurons. In the digging assay, CNO produced a faster onset and robust increase in digging activity in hM3Dq mice, but not in mCherry controls (Fig. 5B). This effect was confirmed in an independent cohort of hM3Dq mice, and no sex differences were observed (Fig. S5A). In the tail suspension test (Fig. 5C), CNO reduced immobility in hM3Dq mice, but not in mCherry controls. This effect was confirmed in an independent cohort of hM3Dq mice, and no sex differences were observed (Fig. S5B). In the elevated plus-maze (Fig. 5D), CNO increased the distance traveled in the closed arms and open proximal arms, reflecting higher locomotion across the maze, in hM3Dq mice, but not mCherry controls. CNO also increased the number of entries into the proximal and distal segments of the open arms, as well as the time spent at the end of the open arms, reflecting increased exploration of the exposed parts of the maze by hM3Dq mice, but not mCherry controls. These effects were generally confirmed in an independent cohort of hM3Dq mice (Fig. S5C), in which CNO increased the distance traveled, number of entries, and time spent in the proximal and distal segments of the open arms. In this cohort, CNO tended to increase the distance traveled in the closed arms, an effect driven by females. In contrast, CNO significantly reduced the time spent in this zone and this effect was more pronounced in males. Females traveled more distance than males in the closed arms. Conversely, males tended to make more entries and to spend more time in the proximal segments of the open arms compared to females. No other sex differences were detected. In the tail pressure test (Fig. 5E), CNO elevated the mechanical nociceptive thresholds of hM3Dq mice, but not mCherry controls. Overall, we found that PSTN *Crh* neuronal hyperactivity is associated with increased digging activity, increased mobility in response to an inescapable stressor (i.e., active coping strategy), increased exploration of innately aversive spaces (i.e., anxiolytic-like effect), and reduced pain sensitivity. Taken together, these phenotypes reflect a state of behavioral disinhibition and are not consistent with hyperkatifeia. To further explore the possibility that PSTN *Crh* neuronal activity could promote alcohol drinking via negative reinforcement, we sought to determine whether the interoceptive and emotional state produced by their chemogenetic activation is aversive. We used two complementary approaches to measure a putative negative hedonic valence. In a paradigm of chemogenetically induced conditioned taste aversion (inspired by ^46^), *Crh*-Cre hM3Dq mice and mCherry controls consumed similar levels of sucrose during conditioning trials and during the preference test, where they both exhibited a strong preference for sucrose over water (Fig. 5F). In a paradigm of chemogenetic self-stimulation (inspired by ^47^), both groups consumed equivalent amounts of fluid regardless of the CNO concentration (Fig. 5G). Altogether, we did not gather evidence of aversion upon activating PSTN *Crh* neurons.

**Figure 5.**
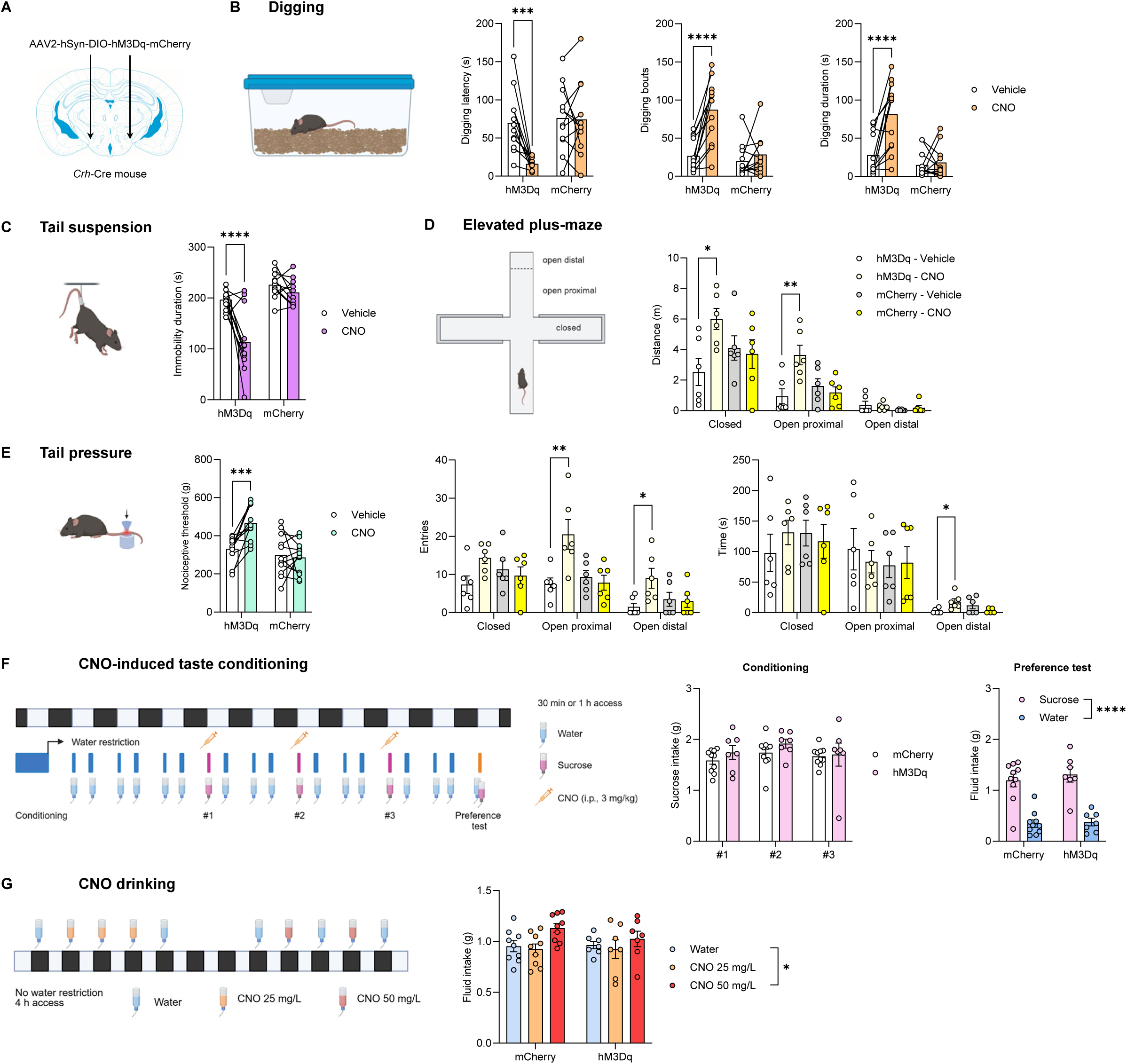
Activating PSTN *Crh* neurons disinhibits behavior in response to novelty and stress. A. *Crh*-Cre mice were injected with a Cre-dependent hM3Dq-encoding vector (or mCherry control) in the PSTN. B-E. CNO increased digging activity (B), mobility in the tail suspension test (C), open arm exploration in the elevated plus-maze (D), and nociceptive threshold in the tail pressure test (E). CNO vs. Vehicle: *, p<0.05; **, p<0.01; ***, p<0.001; ****, p<0.0001, Šídák’s *posthoc* tests. As expected, CNO had no significant effects in mice expressing mCherry only. F. Experimental strategy to measure CNO-induced conditioned taste aversion. Chemogenetic activation of PSTN *Crh* neurons did not affect the amount of sucrose consumed during conditioning trials nor the preference for sucrose over water during 2BC. Main effect of fluid: ****, p<0.0001. G. Mice were given access to regular water or CNO-containing water for 4 h. The mice consumed higher levels of the 50 mg/mL solution, but this effect was not hM3Dq-dependent. Dunnett’s *posthoc* test vs. Water: *, p<0.05. See also Figure S5 and Table S2.

### Escalation of alcohol drinking driven by PSTN activity in dependent mice does not generalize to saccharin

We sought to determine whether the influence of PSTN neurons on alcohol drinking might extend to another fluid reinforcer, saccharin. Mouse cohorts that had been tested for alcohol drinking were subsequently trained to consume saccharin in the same protocol (2-h 2BC).

Stimulating PSTN *Crh* neurons via hM3Dq produced a robust increase in saccharin intake (Fig. 6A-B). To rule out a potential role of testing order, whereby saccharin consumption might have been influenced by the prior consumption of alcohol, a separate cohort of *Crh*-Cre mice expressing hM3Dq in the PSTN was first given access to saccharin. CNO also increased saccharin intake in these mice (Fig. 6C). Divergent effects on alcohol vs. saccharin drinking emerged when probing specific pathways (Fig. 6D) and ensembles (Fig. 6J). Stimulating PVT-projecting PSTN *Crh* neurons, which increased alcohol intake (Fig. 4L), did not affect saccharin drinking (Fig. 6E). Moreover, CeA projectors reduced saccharin intake (Fig. 6F), while rostral VP projectors had no influence (Fig. 6G), even though both populations reduced alcohol consumption (Fig. 4H,J). Likewise, general inhibition of PSTN *Crh* neurons reduced saccharin consumption in both Air and CIE mice (Fig. 6H) but restricting the inhibition to PVT projectors had no impact in either group (Fig. 6I), even though both manipulations reduced alcohol intake (Fig. 2C, 4O). Finally, re-activating the ensembles of PSTN neurons active during early or late withdrawal from CIE reduced saccharin intake in all groups (Fig. 6J-L), again dissociating the influence of endogenous PSTN activity on alcohol vs. saccharin drinking (Fig. 1K vs. Fig. 6J).

**Figure 6.**
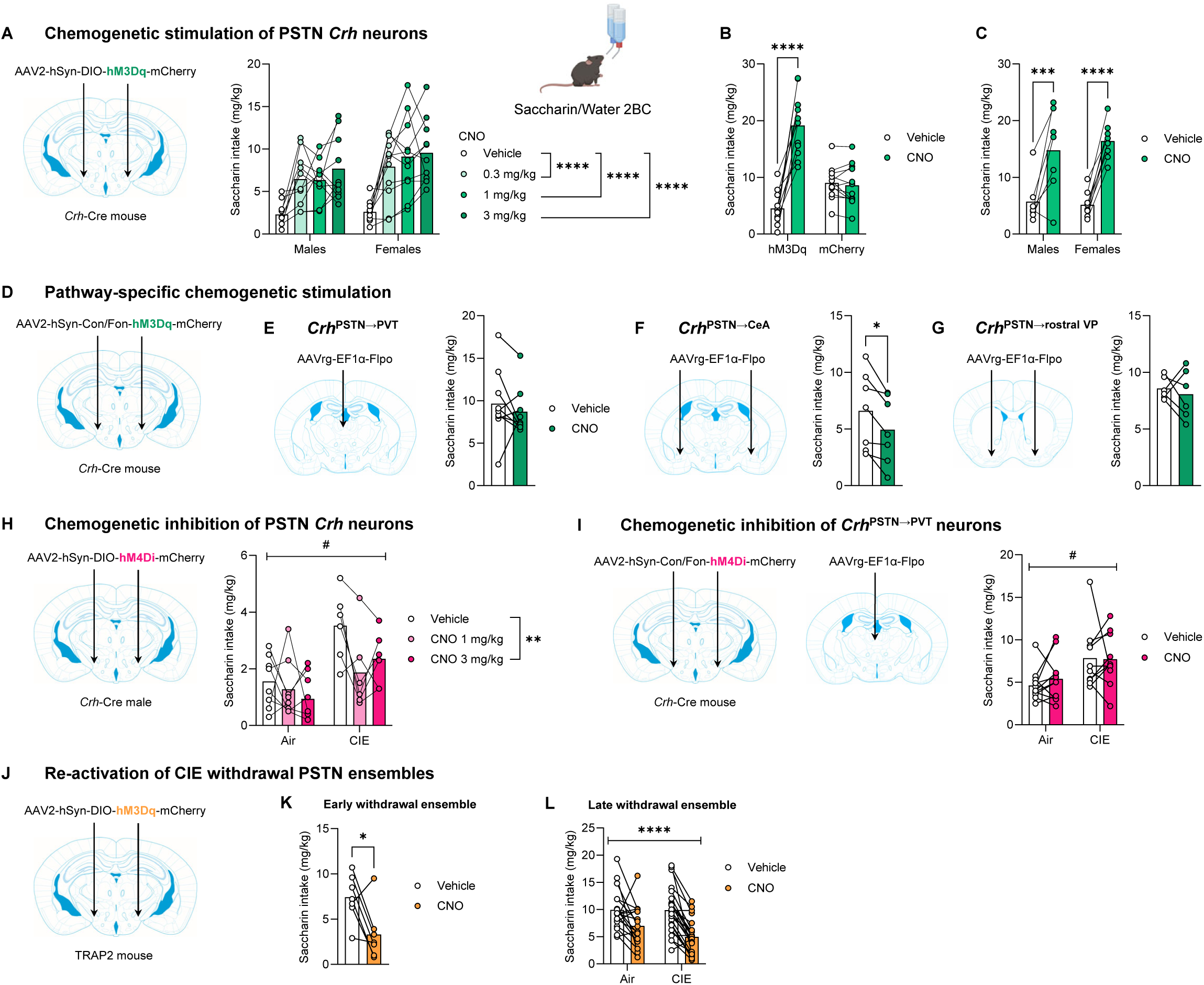
Escalation of alcohol drinking driven by PSTN activity in dependent mice does not generalize to saccharin. A-C. Activating PSTN *Crh* neurons increased saccharin intake in both sexes (A), selectively in hM3Dq-expressing mice (B), and regardless of prior alcohol drinking experience (C). Comparisons to Vehicle: ***, p< 0.001; ****, p<0.0001. D-G. Mice were prepared as in Figure 4G-L, with hM3Dq expression in PSTN *Crh* neurons projecting to the PVT (G), CeA (H), or rostral VP (I). Activating CeA projectors reduced saccharin intake, while activating PVT or rostral VP projectors had no effect. Vehicle vs. CNO: *, p<0.05. H. Inhibiting PSTN *Crh* neurons reduced saccharin intake in Air-2BC and CIE-2BC mice. Main effect of CIE: #, p<0.05. Comparison to Vehicle: **, p<0.01. I. Mice were prepared as in Figure 4M-O, with hM4Di expression in *Crh*^PSTN→PVT^ neurons. CNO did not affect saccharin intake. Main effect of CIE: #, p<0.05. J-L. Mice were prepared as in Figure 1D or 1H, with hM3Dq expression in PSTN cells active during early (K) or late (L) withdrawal from CIE-2BC. CNO reduced saccharin intake in all groups. K. Vehicle vs. CNO: *, p<0.05. L. Main effect of CNO: ****, p<0.0001. See also Table S2.

## Discussion

In this study, we generated multi-pronged evidence that the PSTN undergoes substantial activity changes upon induction of alcohol dependence. Importantly, this reconfiguration directly contributes to alcohol drinking escalation by eliminating the inhibition exerted by PSTN activity in moderate drinkers while recruiting a new PSTN ensemble that promotes alcohol intake. The *Crh* subpopulation participates in this dual influence of the PSTN on alcohol intake through projection-specific effects.

First, we found that PSTN activity strongly increases in early withdrawal from CIE, which transiently suppresses alcohol drinking, but normalizes in late withdrawal, by the time dependent mice show increased motivation for alcohol consumption. Strikingly, despite their similar sizes, the ensembles of PSTN cells active in control and CIE mice at the latter timepoint exert opposite influences on alcohol drinking: PSTN activity suppresses alcohol consumption in non-dependent mice but promotes it in dependent mice. Numerous studies have examined the dynamic patterns of neuronal activation across the brain in response to alcohol consumption, seeking, or withdrawal ^29,48–51^, yet only a handful have investigated the functional impact of these ensembles ^52–57^. To the best of our knowledge, our report is the first to identify a brain region where active ensembles drive opposite effects on alcohol drinking as a function of alcohol dependence. Furthermore, the escalation of alcohol drinking driven by the late withdrawal PSTN ensemble in dependent mice is specific for this substance as the same ensemble reduces saccharin drinking. Such dichotomy positions the late withdrawal PSTN ensemble as a tangible neural substrate for AUD narrowing the focus on alcohol at the expense of alternative, non-drug sources of reward ^58^.

We had hypothesized that the *Crh* subpopulation of PSTN neurons would contribute to alcohol drinking escalation. In support of this hypothesis, we found that the excitability of a subset of PSTN *Crh* neurons is elevated in late withdrawal from CIE and correlates positively with recent alcohol intake, thus providing a novel cellular basis for alcohol drinking escalation in dependence. Furthermore, we demonstrate that stimulating PSTN *Crh* neurons is sufficient to robustly increase voluntary alcohol consumption. This effect does not result from increased thirst but does generalize to saccharin. Conversely, inhibiting PSTN *Crh* neurons reverses the escalation of alcohol and saccharin intake produced by CIE exposure. The PSTN is generally known to suppress consummatory behaviors ^7^. It becomes activated upon sudden food ingestion, exposure to aversive stimuli that reduce feeding (e.g., visceral malaise, novelty, predator odor), and administration of anorectic hormones, thus encoding states of satiety and food rejection. The functional manipulation of PSTN glutamatergic neurons as a whole or subsets of PSTN neurons (e.g., those expressing *Tac1* or *Adcyap1*) demonstrated that their activation serves to suppress food or sucrose intake ^15–21^. Consistent with prior literature, activating PSTN *Tac1* neurons virtually ablated the consumption of alcohol in our experimental conditions. In this context, our finding that PSTN *Crh* neurons promote the consumption of alcohol and saccharin sets these neurons apart as a unique subset opposing the influence of neighboring populations. In a recent study, we found that activating PSTN *Crh* neurons promotes the consumption of a novel, palatable food (Froot Loops) in hungry mice, and the consumption of a novel, palatable fluid (sucrose) in thirsty mice ^21^. Here, we show that this effect extends to alcohol and saccharin drinking, does not require fluid deprivation, and withstands the concomitant availability of water (free-choice consumption) and extensive habituation to the reinforcers. The present study represents the first investigation of the PSTN in relation to psychotropic substances and future studies will determine whether its influence on drug self-administration is specific to orally ingested reinforcers or extends to intravenously infused reinforcers.

In contrast to our prediction, CRF_1_ receptors or other CRF targets do not mediate the increase in alcohol drinking driven by PSTN *Crh* neurons. Testing of alternative signaling mechanisms pointed to a combined role of mGlu5, NK1, and opioid receptors. Each of these receptors has been implicated in the control of alcohol seeking and intake and represents a potential or actual target for the treatment of AUD ^59–61^. Several of the ligands we tested tended to lower alcohol intake in control conditions but became ineffective following CNO administration, suggesting that the pro-drinking subset of PSTN *Crh* neurons shows limited activity under non-dependent conditions. In the presence of the highest dose of mGlu5, NK1 or opioid receptor antagonist, alcohol intake was still 2-3 times higher after CNO than vehicle injection indicating that the contributions of these signaling pathways might be cumulative or that additional pharmacological mechanisms remain to be discovered.

Anterograde viral mapping confirmed that PSTN *Crh* neurons strongly innervate extended amygdala subregions, as previously reported ^17^, but also identified projections to the VP and PVT, two areas that process both rewarding and aversive signals ^62,63^. Pathway-selective manipulations revealed a surprising heterogeneity in the effect of different PSTN *Crh* forebrain projections on alcohol drinking, with CeA and rostral VP projectors reducing intake vs. PVT projectors increasing intake. It is likely that additional projections promoting alcohol intake remain to be identified and that their concurrent activation might be needed to overcome the suppression from CeA and rostral VP projectors and produce the net effect of global PSTN *Crh* neuron stimulation (i.e., robust increase in alcohol drinking). We had previously determined that CeA-projecting PSTN *Crh* neurons accelerate refeeding following food deprivation but do not affect the amount of food or water consumed under states of hunger or thirst, respectively ^21^.

The suppression of alcohol and saccharin drinking reported here therefore seems specific to non-homeostatic consumption. While no previous study had manipulated the *Crh* subset of PVT-projecting PSTN neurons, global activation of the PSTN→PVT pathway in vGluT2-Cre mice suppresses sweet food intake ^15^. Our finding that *Crh*^PSTN→PVT^ neurons promote alcohol drinking and do not influence saccharin consumption thus suggests functional heterogeneity within the PSTN→PVT pathway, consistent with the fact that it also contains a dense *Tac1* component ^17^. PVT activity can be increased by and promote alcohol seeking and drinking ^64^; our data demonstrate that PSTN *Crh* inputs contribute to these effects.

With respect to affect, activating PSTN *Crh* neurons reproducibly increased digging activity (as seen during withdrawal from CIE ^65,66^), mobility in the tail suspension test (as previously reported ^22^), and exploration of the exposed arms of an elevated plus-maze (suggesting an anxiolytic-like effect). It also reduced pain perception in a mechanical nociception assay and did not produce aversion. This combination of phenotypes does not support the notion that PSTN *Crh* neurons would elicit a state of emotional or physical distress (hyperkatifeia) that would promote alcohol consumption via negative reinforcement. Instead, the behavioral changes produced by the stimulation of PSTN *Crh* neurons are consistent with a general pattern of behavioral disinhibition, whereby activity is increased in conditions of high arousal. This pattern is consistent with the effect of stimulating PSTN calretinin neurons (∼90% of PSTN neurons), which increases wakefulness and exploratory behaviors ^67^, but opposite to the influence of BNST-projecting PSTN excitatory neurons, which drive anxiety-like behaviors in response to chronic stress ^27^. In humans, specific facets of behavioral disinhibition and impulsivity are linked to alcohol use via a common externalizing factor ^68–70^. Our results suggest that dysregulation of PSTN *Crh* neurons could represent a shared mechanism driving these traits and the propensity to consume more alcohol. As such, the PSTN might have functional relevance for the “Reward type” neurobehavioral profile of alcohol addiction, which is characterized by higher sensation-seeking and risk-taking, and displays high resting-state connectivity in the salience network ^4^.

The findings reported here may impact future clinical research by drawing attention to the potent and pleiotropic influence of a nucleus that is unbeknownst to most brain experts yet resides in the immediate vicinity (at least in mice, rats, and macaques) of the subthalamic nucleus (STN), a major target of deep brain stimulation in the treatment of Parkinson’s disease and obsessive-compulsive disorder ^26,71,72^. The existence of the PSTN in the human brain remains to be established, but the extensive similarity between the neurochemical features, cortical connectivity, and functional impact of the rodent PSTN and human medial subthalamic region suggests that a homologous nucleus also exists in humans, near the limbic (ventromedial) region of the STN, as discussed in detail by Barbier *et al.* ^73^. Relevant to the proposed link between PSTN activity and reward-driven alcohol addiction, deep brain stimulation of the limbic tip of the STN, which is likely to encroach on the putative PSTN, enhances functional connectivity within the salience network ^74^. Accordingly, the PSTN may represent an unrecognized node of the brain network(s) mediating the dysregulation of incentive salience in AUD ^4,5^, which could shape novel approaches for individualized treatment. Moreover, pre-existing dysfunction of PSTN circuits could form a novel basis for vulnerability to excessive reward seeking. Our findings on the bidirectional influence of specific PSTN cell types and pathways on alcohol drinking also open new avenues for pharmacological modulation of alcohol use in humans. For instance, single-cell molecular profiling of these neuronal subtypes may identify differentially expressed molecules that could be targeted to dampen the recruitment of pro-drinking circuits or re-engage drinking suppression mechanisms weakened in AUD.

## CRediT authorship contribution statement

**JLD:** Conceptualization, Investigation, Analysis, Supervision, Writing - Review & Editing. **MK**: Investigation, Analysis, Validation. **AO:** Investigation, Analysis, Validation. **CL:** Investigation, Analysis, Supervision. **FPV:** Investigation, Analysis, Writing - Review & Editing. **MB:** Investigation, Analysis, Validation, Writing - Review & Editing. **RJS:** Investigation, Resources. **CM:** Investigation. **CK:** Investigation. **LS:** Investigation, Writing - Review & Editing. **AW:** Conceptualization, Writing - Review & Editing. **GM:** Investigation. **HS**: Investigation. **HCB:** Supervision, Resources. **MR:** Project administration, Supervision, Analysis. **KD:** Methodology, Resources. **CC**: Conceptualization, Funding acquisition, Project administration, Supervision, Investigation, Analysis, Visualization, Writing - Original Draft.

## Acknowledgments

This work was supported by National Institutes of Health research grants AA026685, AA027636, AA006420, AA027372, AA030807, AA032709, and AA013498, as well as training grant AA007456 and fellowships AA024952 and AA031926. These funding sources were not involved in study design, data collection, analysis, or interpretation, nor decision to publish. We wish to thank Thao Ngyuen, Sophia Zhu, Tanvi Shah, Ellie Petty, and Maya Mehta for their assistance with histological analysis. We also thank Charu Ramakrishnan and Nadya Andini for preparing the Con/Fon vectors. Some of the schematics were created with BioRender.com.

## Declaration of interests

CC is an employee of Johnson & Johnson. KD is a co-founder and scientific advisor for MapLight Therapeutics and Stellaromics and holds equity in both companies. KD is also a consultant for Modulight.bio and RedTree. These companies did not provide funding for this study and were not involved in design, data collection, analysis, or interpretation, nor decision to publish. Other authors have no competing interests to declare.

## Resource availability

All data supporting the findings of this study are available within the article and its supplementary information file. Raw data will be shared by the lead contact upon request.

## Supplemental figure titles and legends

**Figure S1. The influence of PSTN activity on alcohol drinking switches from inhibitory to stimulatory upon induction of alcohol dependence, related to Figure 1**.

A. In mice exposed to chronic intermittent alcohol inhalation without alcohol drinking, alcohol intoxication reduces, while early withdrawal increases the number of c-Fos+ cells in the PSTN. Air vs. CIE: **, p<0.01.

B. Alcohol-exposed TRAP2 mice injected with a Cre-dependent vector into the PSTN do not express the mCherry reporter in the absence of 4-OHT administration.

C. Blood ethanol concentrations measured during alcohol vapor inhalation in the experiment reported in Figure 1D-G.

D. As in Figure 1F but with CNO dihydrochloride. Pairwise comparisons: *, p<0.05; **, p<0.01; ****, p<0.0001.

E. As in Figure 1I but restricted to males. Air-2BC vs. CIE-2BC: **, p<0.01.

F. Blood ethanol concentrations measured in males during alcohol vapor inhalation in the experiment reported in Figure 1H-L.

G. As in Figure 1J but restricted to males. Pairwise comparisons: *, p<0.05; ****, p<0.0001.

H. As in Figure 1K but restricted to males. Pairwise comparisons: ***, p<0.001; ****, p<0.0001.

I. As in Figure 1I but restricted to females.

J. Blood ethanol concentrations measured in females during alcohol vapor inhalation in the experiment reported in Figure 1H-L.

K. As in Figure 1J but restricted to females. Pairwise comparisons: *, p<0.05.

L. As in Figure 1K but restricted to females. Pairwise comparisons: *, p<0.05; ***, p<0.001; ****, p<0.0001.

See also Table S2.

**Figure S2. Validation of chemogenetic manipulations of PSTN *Crh* neurons and effect of optogenetic stimulation, related to Figure 2**.

A. Blood ethanol concentrations measured during alcohol vapor inhalation in the experiment reported in Figure 2A-L.

B. In slices from mice prepared as in Figure 2A, bath application of CNO reduced the number of spikes evoked by 40-140 pA currents in mCherry-labeled PSTN *Crh* adapting cells. Baseline vs. CNO: *, p<0.05; **, p<0.01; ***, p<0.001; ****, p<0.0001.

C. No change in spike numbers during the same period in the absence of CNO.

D. Immunolabeling of mCherry (red) and c-Fos (green) in the PSTN of mice prepared as in Figure 2M and injected with CNO (1 mg/kg) 90 minutes prior to perfusion. Scale bar, 50 μm.

E-F. CNO increased the number (E) and fraction (F) of cFos+ cells among mCherry+ cells. ***, p<0.001; ****, p<0.0001.

G. Experimental strategy for optogenetic activation of PSTN *Crh* neurons.

H. *Ex vivo*, blue light triggered sustained firing in mCherry+, but not mCherry- cells. Depolarizing the mCherry- cell at −50 mV elicited spontaneous spiking.

I. *In vivo*, blue light increased alcohol intake during 30-min 2BC. Light OFF vs. ON: **, p<0.001.

See also Table S2.

**Figure S3. Baseline alcohol intake prior to ligand testing and repetition of CP376395 experiment, related to Figure 3**.

A. Baseline ethanol intake in subgroups assigned to different doses of CP376395 (see Figure 3A).

B-D. As in Figure 3A but alcohol intake was measured after 1 h and 2 h of alcohol access. CP376395 (20 mg/kg) pretreatment exacerbated the effect of CNO during the second hour. The 2-h cumulative intake data replicated the results reported in Figure 3A. Main effect of CNO or pairwise comparisons: *, p<0.05; **, p<0.01.

E-M. Baseline ethanol intake in subgroups assigned to different doses of riluzole (E, see Figure 3D), MTEP (F, see Figure 3E), JNJ16259685 (G, see Figure 3F), acamprosate (H, see Figure 3G), aprepitant (I, see Figure 3H-I), SB222200 (J, see Figure 3J), naloxone (K, see Figure 3K), SR142948 (L, see Figure 3M), and almorexant (M, see Figure 3N).

See also Table S2.

**Figure S4. Mapping of projections from PSTN *Crh* neurons and chemogenetic inhibition of PVT projectors, related to Figure 4**.

A-C. *Crh*-Cre mice were unilaterally injected in the PSTN with a Cre-dependent vector encoding tdTomato and sypEGFP. tdTomato immunolabeling (red) shows transduced cells and projecting axons, while EGFP immunolabeling (green) shows presynaptic terminals in target areas. In the pictured brain, most transduced cells (>90%) were located within the boundaries of the PSTN; a minority of transduced cells invaded the lateral hypothalamus (LH) at the most anterior level (A), the zona incerta (ZI) at the most dorsal level (B), and the medial terminal nucleus (MT) at the most posterior level (C).

D-R. Innervation was observed in several forebrain, midbrain and brainstem areas.

The field of view is outlined in the brain atlas diagram on the left; distance is anteroposterior from bregma; abbreviations from the Franklin and Paxinos atlas ^75^ are used (CeM, central amygdaloid nucleus, medial division; DR, dorsal raphe nucleus; DRD, DR, dorsal part; EA, extended amygdala; InG, intermediate gray layer of the superior colliculus; IPAC, interstitial nucleus of the posterior limb of the anterior commissure; IRt, intermediate reticular nucleus; LPB, lateral PB; MCLH, magnocellular nucleus of the LH; MPB, medial PB; mRt, mesencephalic reticular formation; PAG, periaqueductal gray; PB, parabrachial nucleus; PBP, parabrachial pigmented nucleus of the VTA; PCRtA, parvicellular reticular nucleus, alpha part; PF, parafascicular thalamic nucleus; PIF, parainterfascicular nucleus of the VTA; PN, paranigral nucleus of the VTA; PSTh, parasubthalamic nucleus; PTg, pedunculotegmental nucleus; PV, paraventricular thalamic nucleus; Re, reuniens thalamic nucleus; ST, bed nucleus of the stria terminalis; STLP, ST, lateral division, posterior part; STLV, ST, lateral division, ventral part; STMAM, ST, medial division, anteromedial part ; STMV, ST, medial division, ventral part; Su3, supraoculomotor PAG; Su5, supratrigeminal nucleus; VDB, nucleus of the vertical limb of the diagonal band; VLPAG, ventrolateral PAG; VP, ventral pallidum; VTA, ventral tegmental area; ZIC, ZI, caudal part; ZIV, ZI, ventral part). Scale bars, 100 μm.

S. As in Figure 4N but restricted to males. Air-2BC vs. CIE-2BC: ***, p<0.001.

T. Blood ethanol concentrations measured in males during alcohol vapor inhalation in the experiment reported in Figure 4M-O.

U. As in Figure 4O but restricted to males. Main effect of vapor: ###, p<0.001. Main effect of CNO: **, p<0.01.

V. As in Figure 4N but restricted to females.

W. Blood ethanol concentrations measured in females during alcohol vapor inhalation in the experiment reported in Figure 4M-O.

X. As in Figure 4O but restricted to females.

See also Table S2.

**Figure S5. Activating PSTN *Crh* neurons disinhibits behavior in response to novelty and stress in both sexes, related to Figure 5**.

A-C. Mice were prepared as in Figure 5A. CNO increased digging activity (A), mobility in the tail suspension test (B), and open arm exploration in the elevated plus-maze (C) in both males and females. Main effect of CNO: *, p<0.05; **, p<0.01; ***, p<0.001; ****, p<0.0001. Main effect of sex: ^#^, p<0.05.

See also Table S2.

## Supplemental information

**Table S1. Details of experimental cohorts.**

**Table S2. Results of statistical analysis.**

## STAR Methods

### Animals

Fos^2A-iCreER^ (TRAP2, stock #030323 ^30^), *Crh*-IRES-Cre (*Crh*-Cre, stock #012704 ^76^), and *Tac1*-IRES2-Cre-D (*Tac1*-Cre, stock #021877 ^77^) breeders were obtained from The Jackson Laboratory. Backcross breeders (C57BL/6J mice from Scripps Research rodent breeding colony) were introduced every 1-2 years to prevent genetic drift. All TRAP2, *Crh*-Cre, and *Tac1*-Cre mice used for experimentation were heterozygous (Het). All experiments included mice from both sexes, except for the CIE-2BC experiment in hM4Di-expressing *Crh*-Cre mice, which used males only based on more robust alcohol intake escalation in this sex ^35–39^.

Mice were maintained on a 12 h/12 h light/dark cycle. Food (Teklad LM-485, Envigo) and reverse osmosis purified water were available *ad libitum* except for a 4-h period of water deprivation prior to water intake measurement (Fig. 2R) and daily water restriction in the CNO-induced taste conditioning experiment (Fig. 5F). Sani-Chips (Envigo) were used for bedding substrate. Mice were at least 10 weeks old at the time of surgery and were individually housed throughout the duration of the experiments, starting at least 3 days prior to behavioral testing. In our experience, single housing reduces interindividual variability in alcohol intake and other behavioral readouts, possibly as a result of ablating social hierarchy and creating consistent levels of social isolation stress in all mice. All tests were conducted during the dark phase under red light unless otherwise specified.

All procedures adhered to the National Institutes of Health Guide for the Care and Use of Laboratory Animals and were approved by the Institutional Animal Care and Use Committee of The Scripps Research Institute.

For cFos immunohistochemistry in C57BL/6J mice exposed to CIE-2BC, immunolabeled sections from a published study were used; all details pertaining to animals, alcohol intake, and immunolabeling protocol are described in that publication ^29^.

### Viral vectors

Details of the viral vectors used in each experimental cohort are provided in Table S1.

Adeno-associated viral (AAV) vectors encoding designer receptors for chemogenetic excitation (hM3Dq) or inhibition (hM4Di, KORD) ^31–33^ fused to the fluorescent reporter mCherry (hM3Dq, hM4Di) or mCitrine (KORD), under the control of the human synapsin promoter and in a Cre-dependent manner, were obtained from Addgene (plasmids #44361, #44362, and #65417, respectively). The hM3Dq and hM4Di constructs were packaged into an AAV2 capsid by the Vector Core at the University of North Carolina (UNC) at Chapel Hill. The KORD construct was packaged into an AAV8 capsid by Addgene.

An AAV8 vector encoding the excitatory opsin humanized channelrhodopsin-2 (hChR2) containing the H134R substitution to achieve higher currents ^78^, fused to mCherry, under the EF1α promoter, and in Cre-dependent manner was obtained from Addgene (plasmid #20297). An AAV2 vector expressing tdTomato and synaptophysin-fused EGFP (separated by a T2A self-cleaving peptide) in a Cre-dependent manner was obtained from UNC Vector Core (Addgene plasmid #51509 ^79^).

AAV8 vectors expressing hM3Dq-mCherry or hM4Di-mCherry under a short EF1α promoter in a Cre- and Flp- dependent manner were generated, packaged, and purified by the laboratory of Karl Deisseroth (AAV8-nEF-Con/Fon-hM3Dq-mCherry, AAV8-nEF-Con/Fon-hM4Di-mCherry) and used in conjunction with a retrograde AAV encoding the Flpo enzyme under an EF1α promoter (AAVrg-EF1a-Flpo), as described in ^21,42^.

AAV vectors encoding a short hairpin RNA (shRNA) targeting the *Crh* mRNA (shCrh, 5’-GCATGGGTGAAGAATACTTCC-3’, selected using BLOCK-iT RNAi designer, loop sequence 5’-TTCAAGAGA-3’) or a control sequence (shControl, 5’-GTACGGTGAGCTGCGTTATCA-3’) under the control of a U6 promoter, along with a GFP reporter driven by a CBA promoter, were produced by Virovek. The shCrh and shControl constructs were packaged in an AAV8.2 capsid, in which the Phospholipase A2 domain encoded by the VP1 Cap gene is replaced with the corresponding domain from AAV2 to optimize endosomal escape ^80,81^.

### Drugs

4-hydroxytamoxifen (4-OHT) was obtained from Hello Bio (HB6040) and dissolved in 200 proof ethanol at 20 mg/mL with 30-min agitation at 37°C. An equal volume of a 1:4 mixture of castor oil (Sigma-Aldrich, 259853) and sunflower seed oil (Sigma-Aldrich, S5007) was mixed in by pipetting, ethanol was evaporated via vacuum centrifugation, and another volume of oil mixture was added to reach a final concentration of 10 mg/mL. 4-OHT was administered i.p. at a dose of 50 mg/kg.

Clozapine-N-oxide (CNO) freebase was obtained from Enzo Life Sciences (BML-NS105-0025) or Hello Bio (HB1807), dissolved in dimethyl sulfoxide (DMSO), and diluted in 0.9% saline for i.p. injection (10 mL/kg body weight) at a dose of 1 mg/kg (unless specified otherwise), 30 min prior to behavioral testing. The vehicle solution contained 0.5% DMSO. CNO dihydrochloride (Hello Bio, HB6149) was dissolved in saline and injected i.p. at a dose of 3 mg/kg (unless specified otherwise). The difference in standard doses between CNO freebase and CNO dihydrochloride results from our empirical determination of the minimum dose needed to reliably elicit a significant increase in ethanol intake in mice expressing hM3Dq in PSTN *Crh* neurons. CNO dihydrochloride was also used to prepare drinking solutions by dissolving in reverse osmosis water at a concentration of 25 or 50 mg/L.

Salvinorin B (SalB) was obtained from Hello Bio (HB4887) and dissolved in DMSO for subcutaneous (s.c.) injection at a dose of 10 mg/kg, 30 min prior to behavioral testing. A volume of 1 mL/kg body weight was injected using a 250-uL Hamilton syringe.

CP376395 hydrochloride (Tocris Bioscience, 3212), riluzole (Tocris Bioscience, 0768), naloxone hydrochloride (MP Biomedicals, 0219024525), and acamprosate calcium (Tocris Bioscience, 3618) were dissolved in saline. MTEP hydrochloride (R&D Biosystems, 2921) was dissolved in

Tween-80 and diluted in water (final Tween-80 concentration: 10%). JNJ16259685 (Tocris Bioscience, 2333) was dissolved in a vehicle made of 10% Captisol (w:v) in water. Aprepitant (Sigma, 1041904) was dissolved in DMSO and diluted in saline (final DMSO concentration: 1%). SR142948 (Tocris Bioscience, 2309) was dissolved in Tween-80 and diluted in saline (final Tween-80 concentration: 0.05%). Almorexant hydrochloride (Selleck Chemicals, S2160) was dissolved in a vehicle made of 20% Captisol (w:v) in water. SB222200 (Tocris Bioscience, 1393) was dissolved in a saline/0.3% Tween-80 vehicle. PA-915 was synthesized as an HCl salt by Miguel Guerrero and Edward Roberts (Scripps Research), as described in ^82^; it was dissolved in DMSO and diluted in saline (final DMSO concentration: 10%). Each of these ligands was administered i.p., immediately prior to CNO/vehicle administration. Doses were selected based on publications reporting significant behavioral effects in mice ^83–97^.

Ethanol was obtained from PHARMCO-AAPER (200 proof for drinking and i.p. injection, 111000200; 95% for vaporization, 111000190). Pyrazole (Sigma-Aldrich, P56607) was dissolved in saline and administered i.p. Saccharin (Sigma-Aldrich, S1002) was dissolved in drinking water at a 0.02% (w:v) concentration.

Chloral hydrate (Sigma-Aldrich, C8383) was dissolved in water at a concentration of 35% (w:v), and injected i.p.

### Experimental cohorts

Cohort details, including sample size by sex and corresponding figure panels, are provided in Table S1.

c-Fos immunohistochemistry was conducted in two cohorts of C57BL/6J males exposed to CIE (or Air) every other week for five weeks, and euthanized at different withdrawal time points (2, 10, 26, and 74 hours or 7 days) to capture neuronal activity happening 2 hours earlier, as described in ^29^. In one cohort, the mice did not undergo behavioral testing. In the other cohort, the mice were also given access to voluntary alcohol drinking (2-h 2BC sessions) prior to CIE exposure and during the intervening weeks (Fig. 1A). For the 7-day withdrawal time point, one subgroup remained abstinent until euthanasia, while another subgroup was given 2BC sessions for 4 days preceding euthanasia.

A cohort of TRAP2 mice was used to manipulate the ensemble of PSTN neurons active in early withdrawal from CIE (Fig. 1D). These mice were given two weeks of alcohol 2BC and split into two groups of equivalent alcohol intake for assignment to the hM3Dq or hM4Di vector. After surgery, the mice were given one more week of 2BC and underwent four rounds of CIE/2BC alternation. After the fifth week of CIE, 4-OHT was administered ∼22 h after alcohol vapor inhalation ended, prior to dark phase onset. The mice were given a week off before 2BC sessions were resumed for 3 weeks and the effect of CNO freebase was tested within-subjects. The effect of CNO dihydrochloride was tested on a subsequent week to verify reproducibility. The mice were then switched to saccharin 2BC for four weeks and the effect of CNO (dihydrochloride) was again tested within-subjects.

Another cohort of TRAP2 mice was used to manipulate the ensemble of PSTN neurons active in late withdrawal from CIE (Fig. 1H). These mice first underwent surgery. After a week off, they were given two weeks of alcohol 2BC and split into two groups of equivalent alcohol intake for assignment to Air and CIE exposure. After five rounds of CIE (or Air)/2BC alternation, 4-OHT was administered 7 days after alcohol vapor inhalation ended and at the time of expected 2BC start (i.e., ∼22 h after the last 2BC session ended). The mice did not receive alcohol that day. They were then exposed to two more rounds of CIE/2BC, and the effect of SalB was tested within-subjects after extensive habituation to vehicle injections (8 times). After three weeks off, 2BC sessions were resumed for two weeks, and the effect of CNO was tested within-subjects. The mice were then switched to saccharin 2BC for three weeks and the effect of CNO was again tested within-subjects.

The effect of chemogenetic inhibition of PSTN *Crh* neurons was tested in two cohorts of *Crh*-Cre mice exposed to CIE-2BC. The mice were given access to 2BC prior to surgery and exposed to five rounds of CIE (or Air)/2BC prior to within-subjects CNO testing. In one cohort, the mice were then switched to saccharin 2BC for two weeks and the effect of CNO was again tested within-subjects.

Seven independent cohorts of *Crh*-Cre mice (Cohorts 1-7) were used to test the effect of chemogenetic activation of PSTN *Crh* neurons on fluid consumption (alcohol [Cohorts 1-3 and 5-6], saccharin [Cohorts 1-3], water [Cohorts 1, 3, and 4]), affect (digging, tail suspension, elevated plus maze [EPM]; Cohorts 1-3), nociception (tail pressure; Cohort 2), and CNO conditioned taste aversion followed by CNO drinking (Cohort 7, experimental timelines shown in Fig. 5F-G, 2-week break between the two procedures). Cohorts 1, 5, and 6 were used to test the ability of ligands (all cohorts) or *Crh* knockdown (Cohort 5) to block the effect of CNO on alcohol drinking. A subset of Cohort 3 was also used to validate chemogenetic activation via mCherry/c-Fos immunolabeling. Cohort 4 was used to measure water intake in the absence of deprivation. Cohorts 1-7 all included mice expressing hM3Dq; Cohorts 2 and 7 also included a control group expressing mCherry only to control for potential off-target effects of CNO; Cohorts 3-7 were injected with a smaller volume of viral vector to minimize viral transduction in adjacent brain regions. The testing order was as follows: Cohort 1 – alcohol, saccharin, water, EPM, ligand testing; Cohort 2 – alcohol, saccharin, digging, tail suspension, EPM, tail pressure; Cohort 3 – water, saccharin, digging, alcohol, tail suspension. These tests were conducted on different weeks. The selection of tests and order of testing were designed to rule out potential carry-over effects and to perform the most stressful tests at the end. In all 2BC assays, mice were given at least 10 baseline sessions with the relevant reinforcer (alcohol or saccharin) before CNO was tested, and vehicle injections were administered for at least 2 consecutive days prior to the first CNO test for habituation purposes. The effect of CNO was tested according to a within-subject design, except for EPM, which was conducted only once in each mouse. Except for CNO dose-responses, mice did not receive CNO more than once in any given week.

For ligand testing in Cohort 1, mice were split into subgroups of equivalent alcohol intake on the two days preceding testing (Mon-Tue, Fig. S3A,E-M) and assigned to a ligand dose (each subgroup contained the same number of males and females); this dose was administered along with vehicle or CNO (within-subject design, counterbalanced order) on two consecutive days (Wed-Thu). The ligands were tested in the following order, with at least one week between ligands: CP376395, riluzole (30 mg/kg was initially included in the dose-response, but not administered beyond the first testing day due to major sedative effects precluding drinking behavior), aprepitant (10 mg/kg on one week, 25 mg/kg on the subsequent week), SR142948, naloxone (0.2 and 1 mg/kg), almorexant, acamprosate, MTEP, JNJ16259685. CP376395 was retested in Cohort 5, but in this case, all mice were exposed to all four combinations of treatment (Vehicle/CP376395 x Vehicle/CNO). PA-915 and naloxone (10 mg/kg) were tested in Cohort 6, and for each ligand, each mouse was exposed to all four combinations of treatment.

Another cohort of *Crh*-Cre mice was used to test the effect of optogenetic stimulation on alcohol drinking.

A cohort of *Tac1*-Cre mice was used to test the effect of chemogenetic activation of PSTN *Tac1* neurons on alcohol and saccharin drinking. A subset of mice was used for *ex vivo* validation by electrophysiology.

The projections of PSTN *Crh* neurons were mapped out by injecting an anterograde viral tracer unilaterally in *Crh*-Cre mice. Brains were collected 3 months later and processed for immunolabeling of tdTomato (cytoplasmic marker) and GFP (presynaptic marker). One brain showing prominent cell body labeling within the PSTN with limited spillover into adjacent regions (<10% labeled somas) was selected for whole-brain immunolabeling and detailed analysis of axonal and terminal labeling across AP levels (+1.5 mm to −5.5 mm from bregma).

To investigate which neuronal pathway might mediate the increase in alcohol drinking driven by PSTN *Crh* neurons, five groups of *Crh*-Cre mice were prepared for selective expression of hM3Dq in PSTN *Crh* neurons projecting to the CeA, BNST, rostral VP, caudal VP, and PVT, respectively. The effect of CNO on alcohol drinking was first tested according to a within-subjects design. When a significant effect of CNO was detected, the mice were then switched to saccharin 2BC and CNO was again tested within-subjects. Another group of *Crh*-Cre mice was prepared for selective expression of hM4Di in PSTN *Crh* neurons projecting to the PVT. These mice were then subjected to five rounds of CIE (or Air)/2BC alternation followed by within-subject testing of CNO on alcohol drinking. Mice were then exposed to two more rounds of CIE (or Air) with alternating saccharin 2BC drinking followed by within-subject testing of CNO on saccharin drinking.

### Stereotaxic surgeries

Mice were anesthetized with isoflurane and placed in a stereotaxic frame (David Kopf Instruments, model 940). Small holes were drilled in the skull (David Kopf Instruments, 1474) and a viral vector was injected bilaterally into the target region (unilaterally for anterograde tracing). The volumes and stereotaxic coordinates used in each cohort are provided in Table S1. Injections were performed using either a dual syringe pump (Harvard Apparatus) controlling the plungers of 10-μL Hamilton syringes connected to 33-gauge single injectors projecting 5 mm beyond a 26-gauge double guide cannula (Plastics One), or a microinjector pump (World Precision Instruments, UMP3T-2) with an attached 10-μL NanoFil syringe (World Precision Instruments) fitted with a 33-gauge NanoFil blunted tip (World Precision Instruments, NF33BL). The vector was infused at a rate of 50-100 nL/min and the injectors were left in place for at least 2 min to minimize backflow. The scalp was sutured using surgical thread. Mice were left undisturbed for at least one week post-surgery, and at least four weeks elapsed until the effect of CNO was tested.

The reason for the large range of viral vector volumes infused in different cohorts (Table S1) is historical. When this project was started, we used a default volume of 1 μL. We then realized that smaller volumes (75-150 nL) were more appropriate to increase the anatomical specificity of viral transduction in the PSTN and minimize cellular damage, without losing transduction efficiency (with titers > 10^13^ vg/mL). We used higher volumes (400-500 nL) for the Con/Fon vector due to its lower titer and 200 nL for the AAVrg vector to favor a larger spread and likelihood of terminal uptake (target regions are larger than the PSTN). We used 200 nL for the ChR2 vector as the anatomical specificity of optogenetic stimulation was further determined by the site of optical fiber implantation.

For optogenetic stimulation, mice were implanted in the PSTN with a dual mirror tip fiberoptic cannula (DFC_200/250-0.66_6mm_GS2.0_MA45_LR, Doric Lenses) at the time of viral vector infusion. The scalp was not sutured. Instead, a headcap was built using C&B Metabond Quick Adhesive Cement System (Parkell), followed by a layer of Tetric N-Flow composite resin (Ivoclar).

### Alcohol drinking

Food pellets were placed in the bedding instead of the food hopper throughout the duration of the drinking experiments. Two-bottle choice (2BC) drinking sessions were conducted Mon-Fri, starting at the beginning of the dark phase and lasting 2 h. During these sessions, the home cage water bottle was replaced with two 50-mL conical tubes fitted with a rubber stopper and sipper tube assembly and filled with water or ethanol 15% (v:v), respectively. The positions of the water and ethanol bottles were alternated every day, and bottles were weighed at the end of each session. Bottles were also placed in an empty cage to generate spill control values that were subtracted from the weights lost in the alcohol and water bottles of experimental cages. Alcohol preference was calculated by dividing the weight of alcohol solution consumed by the total weight of fluids consumed during the session (alcohol + water) and multiplying by 100. In the absence of actual consumption, spill-corrected weights can be negative (they were never lower than 0.05 g) and preference values can thus be slightly higher than 100%. Body weights were measured on a weekly basis to calculate ethanol intake (g ethanol per kg body weight).

Modified cages were used for alcohol drinking during optogenetic stimulation. These cages featured two holes in one of the shorter walls to insert the conical tubes from the side of the cage, thus allowing removal of the cage lid and unrestricted motion of the mouse while tethered to the patchcord (Doric Lenses, SBP(2)_200/230/900-0.57_2m_FCM-GS2.0), itself connected to a rotary joint (Doric Lenses, FRJ_1x1_PT-400/430/LWMJ-0.57_1.0m-FCM_0.15m-FCM). Mice were trained to drink alcohol in these cages for 30-min sessions and extensively habituated to patchcord tethering prior to testing the effect of optogenetic stimulation (10-ms pulses delivered at 20 Hz with a power of 8 mW). “Light OFF” testing was conducted while the mice were tethered but no light was delivered.

### Alcohol vapor inhalation

Alcohol intake escalation was induced by alternating weeks of voluntary alcohol drinking during limited-access 2BC sessions (described above) with weeks of forced chronic intermittent alcohol exposure via vapor inhalation (CIE) ^28^. During CIE weeks, CIE-2BC mice were exposed to 4 cycles (Mon-Fri) of 16 h ethanol vapor inhalation/8-h air inhalation followed by 72 h withdrawal (Fri-Mon). Ethanol was dripped into a heated flask using a metering pump (Walchem, EWN-B11PEUR), and an air pump (Hakko, HK-40LP) conveyed vaporized ethanol into custom chambers (modified from Allentown Sealed Positive Pressure individually ventilated cages). Mice received an i.p. injection of ethanol (1.5 g/kg) and pyrazole (68 mg/kg) before each 16-h ethanol vapor inhalation session. Blood alcohol levels (BALs) were measured on a weekly basis using gas chromatography and flame ionization detection (Agilent 7820A). The drip rate was adjusted to yield target BALs of 150-250 mg/dL. Control mice (Air-2BC) breathed air only and received pyrazole.

### Saccharin drinking

Saccharin (0.02% w:v) 2BC drinking sessions were conducted as described above for alcohol. Saccharin intake is expressed as mg saccharin per kg body weight.

### Water intake

In one experiment (Fig. 2R), the home cage water bottle was removed for the first 4 hours of the dark phase to stimulate higher levels of water consumption during the test. A single bottle of water (same drinking tube design as described above for 2BC) was provided during the 2-h session. In another experiment (Fig. 2S), the home cage water bottle was replaced with a drinking tube filled with water for 2-h starting at dark onset (i.e., no water deprivation, quenched state).

### Digging test

Digging activity was assessed according to Deacon ^98^. Digging behavior is sensitive to multiple psychotropic drugs (see ^98^ and references therein) and we previously reported that it is robustly increased during withdrawal from CIE ^65^. The mouse was placed in a clean cage with a 5-cm thick layer of bedding (no lid) and allowed to freely dig for 5 min (Fig. 5B) or 3 min (Fig. S5A). The latency to dig, number of digging bouts, and total digging duration were recorded. Testing was conducted under dim white light.

### Tail suspension test

This test was used to assess the coping strategy used by mice facing an acute, inescapable stressor ^99,100^. Mobility is thought to represent an active coping strategy in response to an acute unescapable stressor, to be contrasted with the passive coping reflected by immobility ^101–103^. Alternatively, increased mobility can be interpreted as lack of adaptive learning, whereby switching to immobility favors energy conservation and survival ^104^. The mice were suspended by their tails using adhesive tape wrapped around the tail approximately 2 cm from the tip and affixed to shelving. Prior to taping, the tail was inserted in a clear hollow cylinder (3.5-cm length, 1-cm diameter, 1 g) to prevent tail climbing behavior. The test lasted 6 min. The total duration of immobility was recorded.

### Elevated plus-maze

The apparatus consisted of two opposite open arms (30 cm length × 5 cm width), with a 0.3 cm lip, and two enclosed arms of the same size, with 15 cm high walls. The runways were made of gray (Fig. 5D) or black (Fig. S5C) acrylic and elevated 30 cm above the ground. The lips and walls were made of translucent acrylic. The end of the open arms (starting 5 cm away from the edge) was defined as the distal zone (the proximal zone represents the remainder). Testing began by placing an animal on the central platform of the maze facing an open arm. The test lasted 5 min. The maze was cleaned between subjects. The following measures were recorded using the ANY-maze (Stoelting Co.) behavioral tracking system: total distance traveled, time spent and number of entries in the closed arms, open arms proximal zone, and open arms distal zone. The total distance traveled was used as an index of locomotor activity and the time spent in the open arms was used as an index of anxiety-like behavior, as previously described ^105,106^. The apparatus design (open arms with ledges and closed arms with transparent walls) was meant to encourage exploration of the open arms and facilitate the detection of an anxiogenic-like effect ^107,108^.

### Tail pressure test

Mechanical nociceptive thresholds were assessed by applying pressure on the tail using a digital Randall-Selitto apparatus (Harvard Apparatus), as previously described by Elhabazi and colleagues ^109^. The mice were first habituated to enter a restrainer pouch made of woven wire (stainless steel 304L 200 mesh, Shanghai YiKai) over three days. On testing days, the mouse was gently introduced into the restrainer, and the distal portion of the tail was positioned under the conical tip of the apparatus. The foot switch was then depressed to apply uniformly increasing pressure onto the tail until the first nociceptive response (struggling or squeaking) occurred. The nociceptive threshold, i.e., the force (in g) eliciting the nociceptive response, was recorded. A cutoff force of 600 g was enforced to prevent tissue damage. The measure was repeated on the medial and proximal parts of the tail of the same mouse, with at least 30 s between each measure. The average of the three measures (distal, medial, proximal) was used for statistical analysis.

### CNO-induced taste conditioning

We adapted the procedure described by Chen and colleagues ^46^ to measure the aversive properties of chemogenetic activation using taste conditioning. Mice were individually housed and provided with two bottles (conical tubes) filled with water. After 7 days of acclimation, mice were water-deprived overnight. On days 1-3, mice were habituated to water restriction; they were only given access to water (single bottle) for 30 min approximately 4 h after the onset of the light cycle and for 1 h before the onset of the dark cycle. On days 4, 6 and 8, mice were given 30 min access to a novel 5% sucrose solution (single bottle), followed by an injection of CNO (3 mg/kg, freebase, i.p.); water was available for 1 h in the afternoon. On days 5, 7 and 9, water was provided following the same schedule as days 1-3. On day 10, mice were given access to two bottles (water and 5% sucrose) for 30 min. The volume consumed was recorded in grams and converted to milliliters. Conditioned taste aversion is reflected by a lower sucrose intake in hM3Dq mice compared to mCherry mice, as seen upon stimulation of parabrachial *Calca* neurons ^46^.

### CNO drinking

We adapted the procedure described by Giardino and colleagues ^47^ to measure the rewarding or aversive properties of chemogenetic activation using self-stimulation. We used a single-bottle rather than two-bottle design, as we reasoned that the subjective effects of chemogenetic activation would be separated from ingestion by several minutes ^110^, preventing the mice from differentiating the water bottle from the CNO bottle based on their respective hedonic valence. Mice were individually housed. The home cage water bottle was replaced with a drinking tube containing water or CNO for 4 h each day, starting 3 h into the dark phase. The volume consumed was recorded in grams and corrected for spilling (i.e., the volume lost in an empty cage was subtracted). The fluids were presented in the following order over 2 consecutive weeks: water (1 day), CNO 25 mg/L (3 consecutive days), water (2 days), CNO 50 mg/L (1 day), water (1 day), CNO 50 mg/L (1 day), water (1 day). For statistical analysis, the intake of each fluid was averaged across all the days it was offered. A rewarding effect is reflected by higher consumption of CNO solution in hM3Dq mice compared to mCherry mice, as seen upon stimulation of BNST *Cck* neurons, while an aversive effect is reflected by lower consumption of CNO solution in hM3Dq mice compared to mCherry mice, as seen upon stimulation of BNST *Crh* neurons ^47^.

### Histology

At the end of all experiments, brains were analyzed to evaluate stereotaxic targeting accuracy by visualizing the mCherry and GFP reporters using native fluorescence or immunolabeling. Mistargeted mice (i.e., lacking viral transduction within the PSTN) were excluded from behavioral datasets accordingly (sample sizes reported in Table S1 include well-targeted mice only).

For native fluorescence and immunolabeling, the mice were anesthetized with chloral hydrate and perfused with cold PBS followed by 3.7% paraformaldehyde (PFA). Brains were dissected and immersion fixed in PFA for 2 hours at 4°C, cryoprotected in 30% sucrose in PBS at 4°C until brains sank, flash frozen in isopentane chilled on a dry ice ethanol slurry and stored at −80°C. Coronal 35-µm thick brain sections were sliced with a cryostat (Leica CM1950), collected in five series spanning the PSTN in PBS containing 0.01% sodium azide, and stored at 4°C.

For native fluorescence, sections were washed in PBS, plated on Superfrost plus glass slides (Fisher Scientific, 1255015), and air-dried. Coverslips were mounted using DAPI-containing Vectashield Hardset medium (Vector Laboratories, H1500). Images were captured using a Keyence BZ-X700 fluorescence microscope.

For immunolabeling, the sections were first blocked in PBS containing 0.3% Triton-X100, 1 mg/mL BSA, and 5% normal goat serum (NGS) for 1 h, then incubated with the primary antibody diluted in PBS containing 0.5% Tween-20 and 5% NGS (rabbit anti-mCherry antibody, Abcam, ab167453, RRID:AB_2571870, 1:5,000; chicken anti-mCherry antibody, Abcam, ab205402, RRID AB_2722769, 1:5,000; chicken anti-GFP antibody, Abcam, ab13970, RRID:AB_300798, 1:2000; rabbit anti-c-Fos antibody, Synaptic Systems, 226 008, RRID:AB_2891278, 1:1000) overnight (or 72 h for c-Fos/mCherry double immunolabeling) at 4°C. Following washes in PBS, sections were incubated with the secondary antibody diluted in PBS (goat anti-rabbit conjugated to Alexa Fluor 568, Life Technologies, A11004, RRID:AB_2534072, 1:500; goat anti-chicken conjugated to Alexa Fluor 488, Life Technologies, A11039, RRID:AB_142924, 1:500; goat anti-chicken conjugated to Alexa Fluor 568, Life Technologies, A11041, RRID AB_2534098, 1:500) for 2 h at room temperature, washed in PBS, and mounted and imaged as described above.

To quantify c-Fos induction following chemogenetic activation of PSTN *Crh* cells, *Crh*-Cre mice expressing hM3Dq were injected with vehicle or CNO (freebase, 1 mg/kg, i.p.) 90 min prior to perfusion. Sections were immunolabeled for c-Fos (green) and mCherry (red). Single- and double-labeled PSTN cells were counted in the merged image using the Cell Counter plugin of Fiji ^111^. Counts from both hemispheres were averaged for each mouse.

The co-localization of *Crh* with other genes was assessed in PSTN sections from naïve C57BL/6J mice using the RNAscope Fluorescent Multiplex manual assay (ACD, 320851). Mice were perfused with cold PBS, quickly decapitated, and brains were snap-frozen in isopentane. Ten series of 20-μm coronal sections were sliced in a cryostat, directly mounted on Superfrost slides, and stored at −80°C. The kit protocol was followed (ACD documents 320513 and 320293), except that Protease III was used in lieu of Protease IV, slides were covered with Rinzl plastic coverslips during incubation steps, and probes were hybridized for 3 h. The following probes were used: mouse *Crh* (316091-C2), mouse *Gad2* (439371), mouse *Slc17a6* (319171-C3), mouse *Penk* (318761), mouse *Tac1* (410351), and mouse *Nts* (420441). Sections containing the PSTN were imaged using a Zeiss Axiophot microscope equipped with a QImaging Retiga 2000R color digital camera and QCapture software. PSTN cells containing signal for each probe were counted and the percentage of colocalization with *Crh* was calculated.

### Electrophysiological recordings

Mice were anesthetized (5% isoflurane) and the brains were quickly removed and placed in oxygenated (95% O_2_/5% CO_2_), ice-cold high-sucrose cutting solution (pH 7.3-7.4) containing (in mM): 206.0 sucrose; 2.5 KCl; 0.5 CaCl_2_; 7.0 MgCl_2_; 1.2 NaH_2_PO_4_; 26.0 NaHCO_3_; 5.0 glucose; 5.0 HEPES, as reported ^112^. A Leica VT1200S vibratome was used to cut 300 µm coronal slices.

The slices recovered in oxygenated artificial cerebrospinal fluid (ACSF; in mM): 130.0 NaCl; 3.5 KCl; 2.0 CaCl_2_; 1.25 NaH_2_PO_4_; 1.5 MgSO_4_; 24.0 NaHCO_3_; 10.0 glucose at 32 °C for 30 min and at room temperature for at least 30 min before recordings.

We visualized PSTN neurons with infrared differential interference contrast (IR-DIC) optics, a 60x water immersion objective (Olympus BX51WI), and a CCD camera (EXi Aqua, QImaging). Neurons expressing mCherry were identified using LED fluorescence illumination (Prior Scientific).

Whole-cell patch clamp recordings were conducted using a Multiclamp 700B amplifier, Digidata 1440A digitizer, and pClamp 10 software (Molecular Devices) and borosilicate patch-clamp electrodes with a tip resistance of 3.5-6 MΩ (Warner Instruments). The electrodes were pulled using a PC-10 puller (Narishige International USA) and were filled with internal solution (in mM): K-Gluconate 145; 0.5 EGTA; 2 MgCl_2_; 10 HEPES; 2.0 Na^+^-ATP; 0.2 Na^+^-GTP (adjusted with 1 M KOH to pH 7.2–7.4 and final osmolarity 290–305 mOsm). All recordings were performed at room temperature.

Membrane properties of PSTN neurons clamped at −70 mV in voltage-clamp configuration were determined using pClamp’s “membrane test”. Recordings with access resistance Ra ≥ 20 MΏ were excluded from analysis. An episodic step current-voltage protocol (I-V) was used in the current-clamp mode to record from PSTN neurons held at Vm=-70 mV. The current-clamp recordings were filtered with 10 kHz low-pass filter. The I-V protocol comprised 22 sweeps (individual sweep length 1.3 s) with superimposed 800-ms hyperpolarizing and depolarizing square current steps (Δ=20 pA), starting from −120 pA. The NeuroExpress software (Ver. 24.a.31; developed and kindly provided by Dr. A. Szucs ^113^) was used to determine intrinsic membrane and action potentials (AP) properties. In a subset of cells, CNO was bath applied at a concentration of 200 nM and the I-V protocol was repeated 10 min later, as in ^114^. Rundown control recordings (i.e., repeated I-V protocol in the absence of CNO) were also conducted.

For optogenetic stimulation, whole-cell patch clamp recordings were conducted in current clamp configuration and 10-ms pulses of blue light were applied at different frequencies (1-20 Hz) for 10 sec to verify excitation, as in ^115^.

### Statistical analysis

Data analysis was performed in GraphPad Prism (v10.2.3). The results of all statistical analyses are reported in Table S2. The effects of vapor or CNO on c-Fos cell counts were analyzed by two-way analysis of variance (ANOVA) or unpaired t-tests. The within-subjects effects of CNO on behavioral measures were analyzed by paired t-test or by repeated-measures (RM) two-way ANOVA with sex, vector, ligand dose, or vapor exposure as between-subjects factor. The effect of optogenetic stimulation was analyzed by paired t-test. Significant interactions were followed by Dunnett’s multiple comparisons to Vehicle for dose-responses, and Šídák’s multiple comparisons otherwise. Select pairwise comparisons (e.g., escalation of ethanol intake on last week of 2BC prior to 4-OHT administration or brain collection) used uncorrected Fisher’s LSD tests. Baseline alcohol intake prior to ligand dose assignment was analyzed by one-way ANOVA or unpaired t-tests. EPM data from each zone were analyzed by two-way ANOVA, followed when relevant by Šídák’s multiple comparisons. All t-tests were two-tailed. For RM ANOVAs, the Geisser-Greenhouse correction was used. Mice were excluded from a given dataset if their ethanol (saccharin, respectively) intake in the vehicle condition was lower than 0.3 g/kg (0.3 mg/kg, respectively), or if their value met with the Grubbs’ outlier criterion (no more than 1 mouse excluded per experimental subgroup) ^116^. In graphs, individual values and group averages are plotted and the error bars represent the standard error of the mean.

