## Supplementary figures and images for "Functional remodeling of the parasubthalamic nucleus drives alcohol drinking escalation in dependence"

### Supplementary Figures

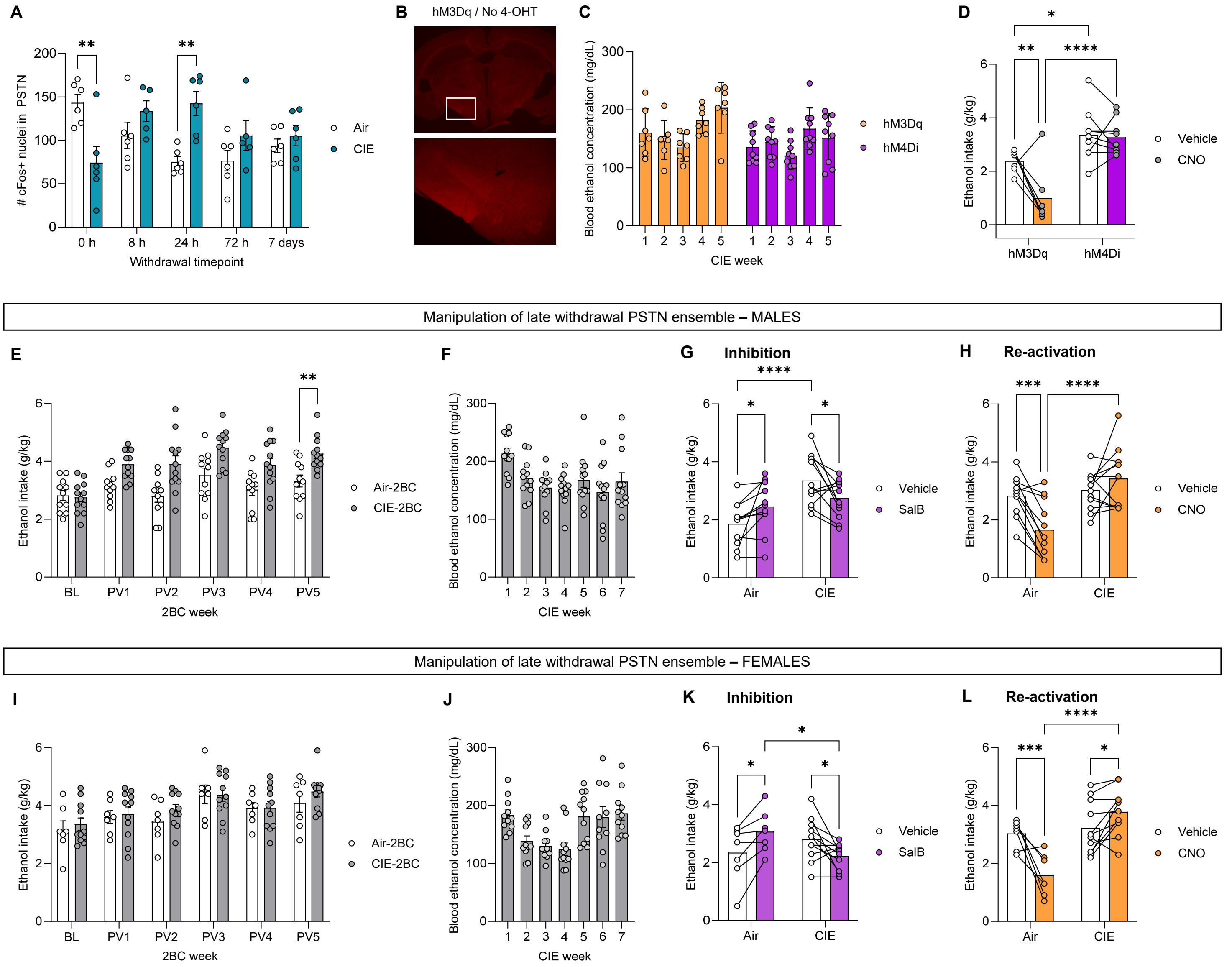

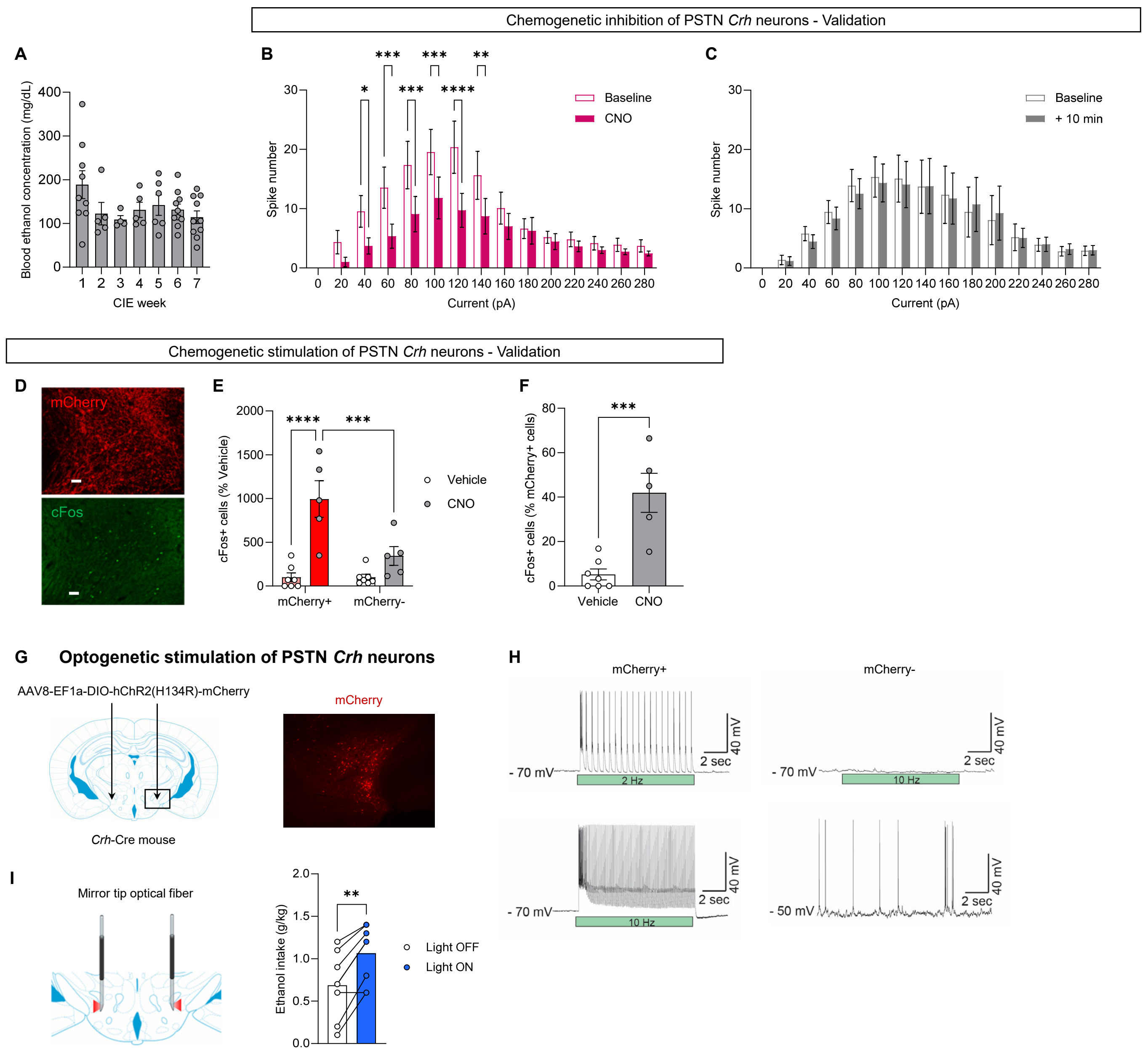

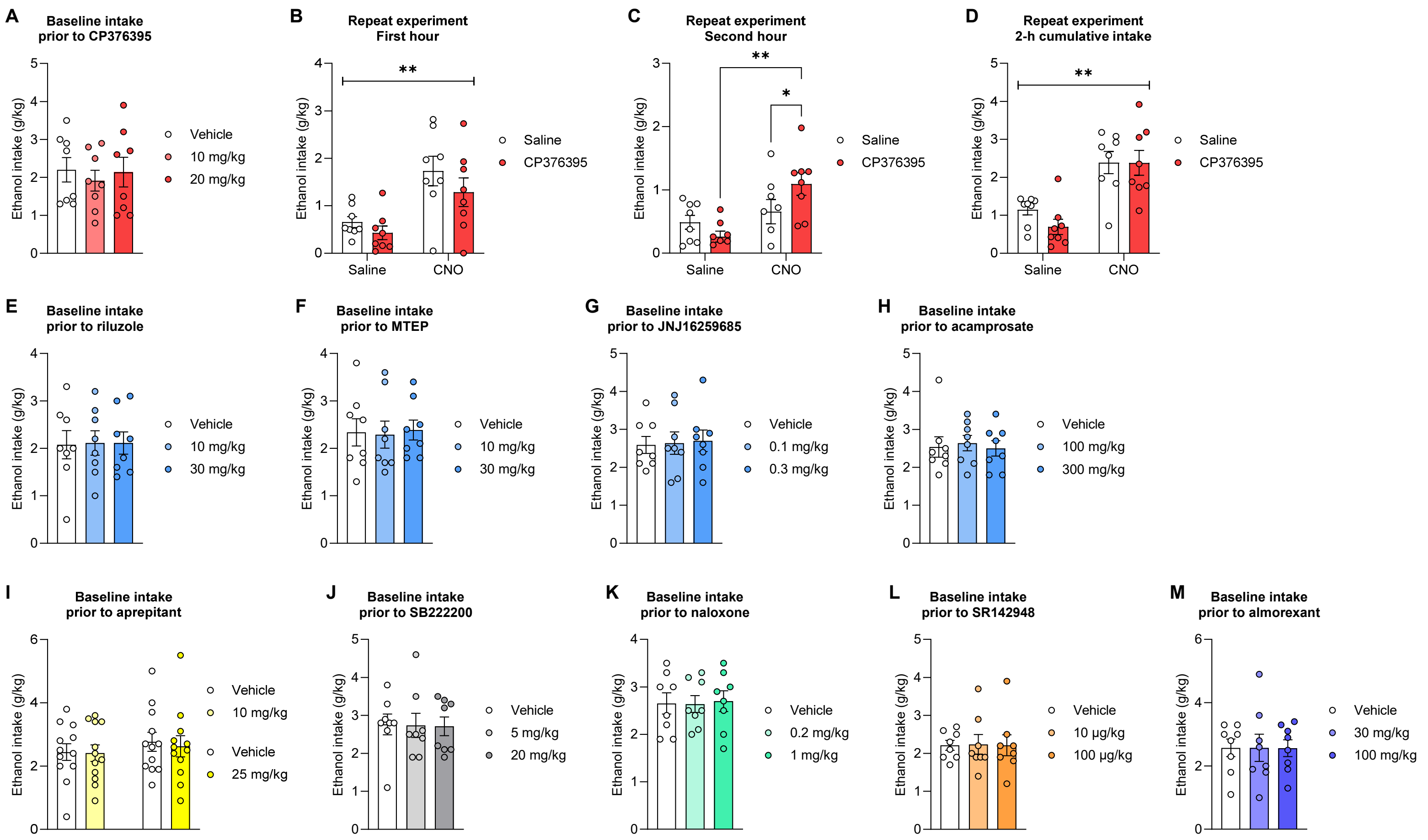

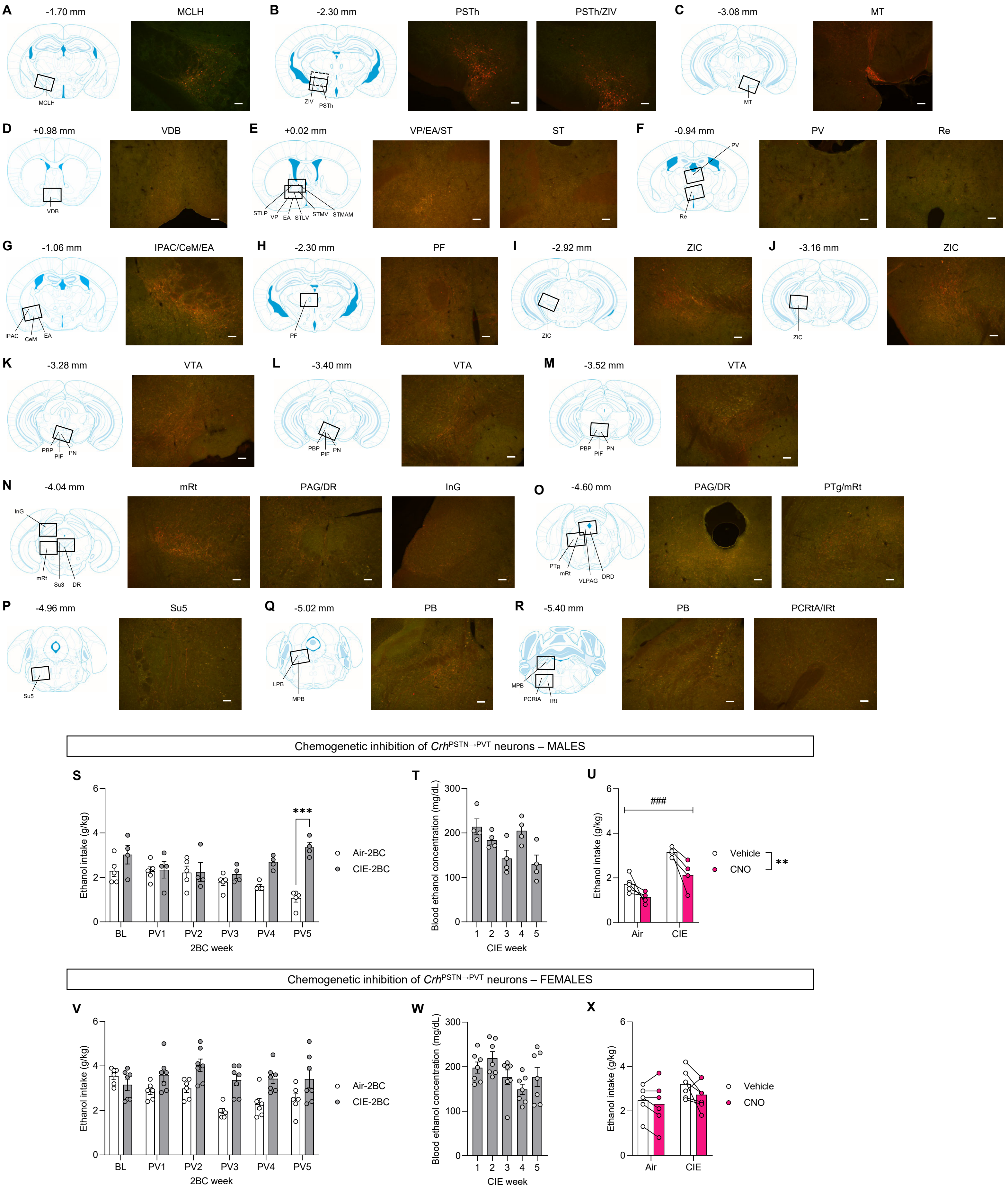

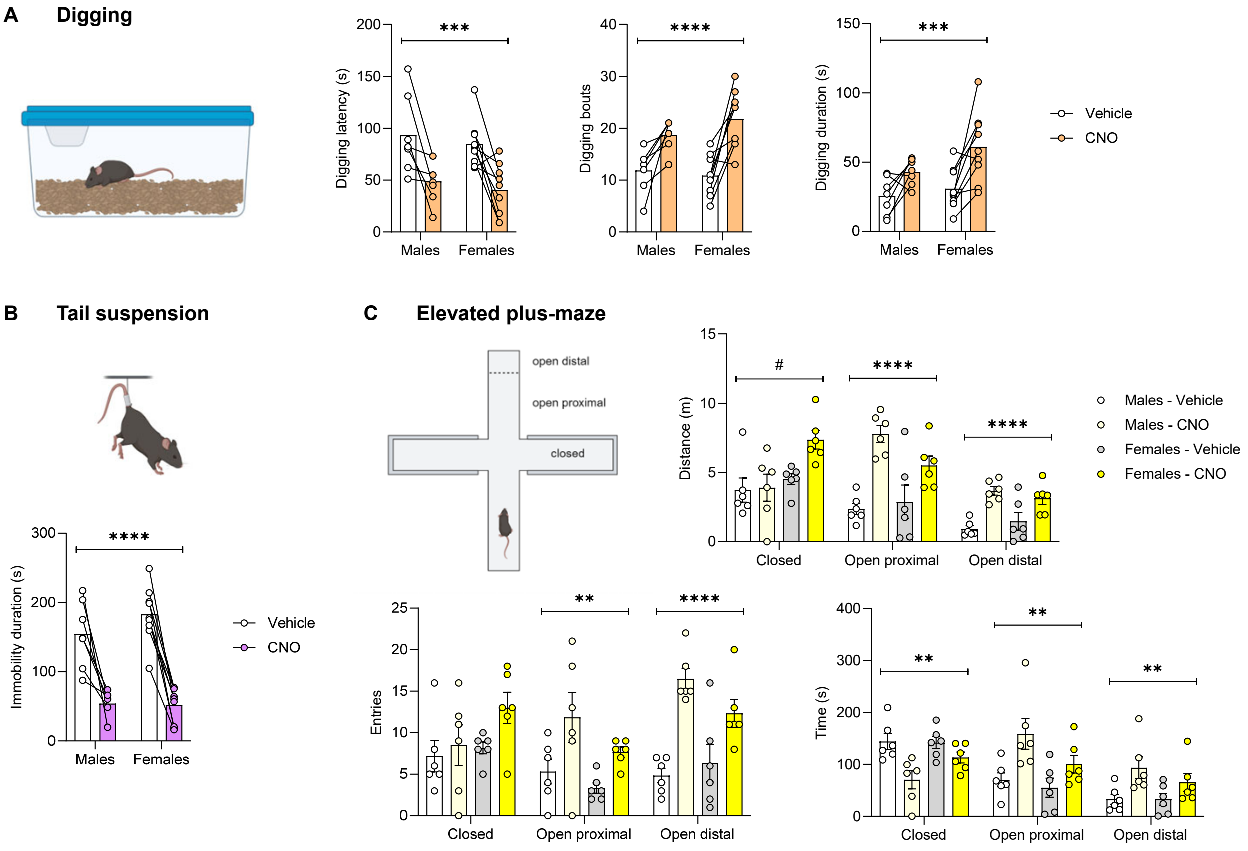
